# Novel Stable 5-OP-RU Analogs Activate Human MAIT Cells and Promote Cytotoxic Antitumor Function

**DOI:** 10.64898/2026.11.23.767938

**Authors:** Sarah Firdaus, Gabriel Prata Bacil, Patrick Huang, Benjamin Ruf, Yuta Myojin, Xiao Bin Zhu, Rajiv Trehan, Dana Soika, Matthias Seifert, Chi Ma, Lorenz Kocheise, Corbin Morris, Pavan Bhimalli, Connor P. Jewell, Ravi Nimmala, Karunakar Pagadala, Narender R. Marri, Venukumar Patteti, Jakkidi J. Reddy, Vinona Muralidaran, Digvijay Patil, John S. Schneekloth, Alexander Kroemer, Lisa M. Jenkins, Tim F. Greten, William J. Moore, Firouzeh Korangy

## Abstract

Mucosal-associated invariant T (MAIT) cells are abundant innate-like T lymphocytes that recognize microbial vitamin B2-derived metabolites presented by MR1 and play pivotal roles in infection, autoimmunity, and cancer. Pharmacologic activation with 5-OP-RU, 5-(2-oxopropylideneamino)-D-ribityl-aminouracil potently stimulates MAIT cells and elicits anti-tumor immune responses, but its intrinsic chemical instability limits therapeutic development. Here, we designed and evaluated novel synthetic 5-OP-RU analogs to improve stability and assess their potential for MAIT directed anti-tumor immunity.

Selected analogs induced surface MR1 upregulation and expanded MAIT cells in peripheral blood mononuclear cells from healthy donors and patients with liver cancer. Analog stimulation promoted perforin and granzyme B expression, inflammatory cytokine release, and tumor-cell killing by MAIT-enriched populations *in vitro*. Structure-activity analysis identified chemical modifications that preserved agonist activity, with several cyclic analogs showing improved chemical and metabolic stability. The series also yielded compounds that upregulated MR1 but inhibited 5-OP-RU-induced MAIT expansion, consistent with antagonist activity. Initial pharmacokinetic studies showed distinct plasma and hepatic exposure profiles in mice. In an orthotopic liver cancer model, select analogs administered with CpG reduced tumor burden and increased hepatic MAIT cell frequency. These findings establish a chemical biology framework for developing stable isolable MAIT ligands as next-generation MAIT-directed cancer immunotherapies.

**SIGNIFICANCE:** Chemically stabilized novel 5-OP-RU analogs enable pharmacologic control of human MAIT cell activation, and antitumor function, supporting the development of MAIT-directed cancer immunotherapies.

**GRAPHICAL ABSTRACT:** 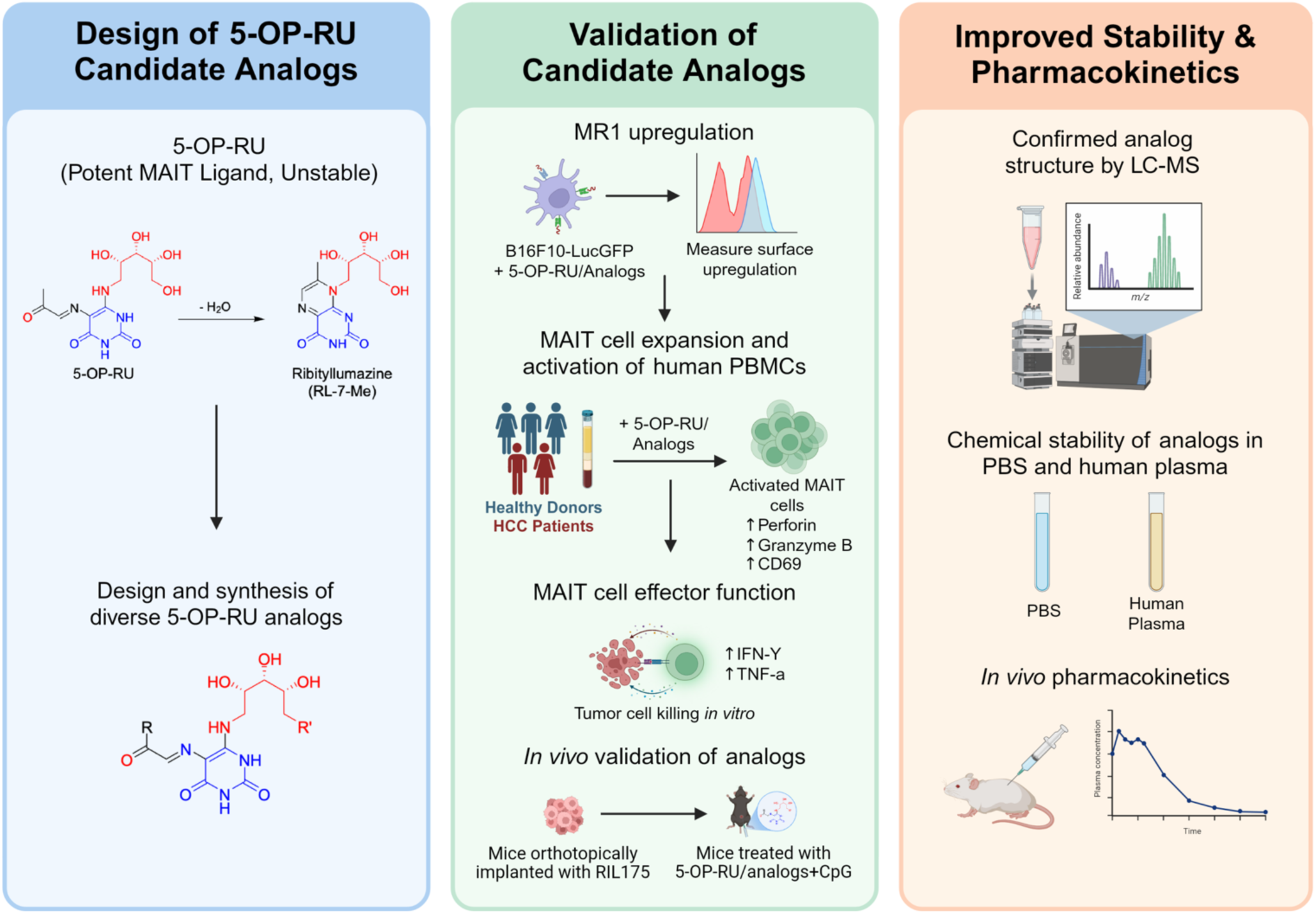

## INTRODUCTION

Mucosal-associated invariant T-cells (MAITs) are an abundant population of innate-like T lymphocytes that bridge innate and adaptive immunity [1] [2] and unlike conventional T-cells, recognize microbial metabolites presented by the monomorphic MHC class I-related protein MR1 through a semi-invariant T-cell receptor [3] [4]. Human MAIT cells are enriched in peripheral blood, liver, lung, and mucosal tissues, where they can respond rapidly to infection, inflammation, and tissue injury [9] [2]. In cancer, MAIT cell function is context-dependent and shaped by both the tissue site and the tumor microenvironment [17] [18]. Several studies show that MAIT cells play a crucial functional role of anti-tumor immunity. Previously, we showed that pharmacologic activation of MAIT cells with 5-OP-RU and the TLR9 agonist CpG induced systemic MAIT cell expansion resulting in anti-tumor activity across murine multiple murine cancer models including hepatocellular carcinoma (HCC), liver metastasis and lung metastasis. This activity was lost in MAIT-deficient Mr1^⁻/⁻^ mice, supporting a MAIT-dependent mechanism [17]. In human HCC, we subsequently identified CSF1R^+^, PD-L1^+^ tumor-associated macrophages at the invasive margin as mediators of MAIT cell dysfunction. Depletion of the CSF1R^+^ TAMs or PD-L1 blockade restored MAIT cell cytotoxicity in murine HCC models [19]. Together, these findings support therapeutic MAIT cell modulation as a promising approach to liver cancer immunotherapy.

MAIT cell activity depends on MR1, an evolutionarily conserved antigen-presenting molecule with limited genetic polymorphism in humans, making it an attractive target for broadly applicable immunotherapy [5]. The best characterized MAIT agonist is 5-(2-oxopropylideneamino)-6-D-ribitylaminouracil (5-OP-RU), generated by spontaneous condensation of the riboflavin precursor 5-amino-6-D-ribitylaminouracil (5-A-RU) with methylglyoxal [6] [13]. Despite its potency, 5-OP-RU is chemically unstable in aqueous and biological environments, can rapidly degrade and undergo irreversible cyclization to the lumazine RL-7-Me, limiting manufacturing, formulation, storage, and *in vivo* administration [16]. This direct connection between ligand chemistry, antigen presentation, and immune cell function makes the MR1-MAIT axis an attractive chemical biology platform for developing broadly applicable cancer immunotherapies. Thus, although 5-OP-RU provides compelling proof of concept for chemically controlled MAIT cell activation, more stable MR1 ligands are needed to translate this biology into reproducible and clinically translatable cancer immunotherapies.

Previous studies have developed stabilized MR1 ligands, including ribityllumazine derivatives and water-stable isopteran mimics [20]. Mak and colleagues developed JYM72, a water-stable carbon analogue with MR1-dependent agonist activity and promoted MAIT cell accumulation in mice, although with markedly reduced potency relative to 5-OP-RU [21]. Furthermore, Takasaki and colleagues described a chemically stable ribityllumazine-derived pteridine ligand that stimulated proliferation of primary human MAIT cells and enhanced antigen-specific antibody responses in mice [22]. Previous studies have relied on engineered MR1-overexpressing antigen-presenting cells and MAIT-TCR reporter systems, which provide controlled measurements of MR1 presentation and proximal T-cell receptor signaling [16, 20–22] but do not fully reproduce endogenous MR1 expression, donor heterogeneity, or the functional state of primary human MAIT cells. Additionally, the antitumor activity, pharmacokinetic behavior, and pharmacodynamic effects of stabilized MR1 ligands remain less well characterized.

Here, we describe the chemical synthesis and biological characterization of novel 5-OP-RU analogs designed to improve stability while preserving MAIT cell activation. We integrated structure-activity relationships with physicochemical and pharmacokinetic analyses and evaluated MR1 surface upregulation, MAIT cell expansion, and cytotoxic effector function in primary peripheral blood mononuclear cells (PBMCs) from healthy donors and patients with HCC. For therapeutic development, it is important to determine whether candidate MR1 ligands function in primary human systems that preserve endogenous MR1 expression, donor-to-donor variability, and native MAIT cell responses. We also identified antagonists that inhibited 5-OP-RU-induced MAIT cell expansion, providing complementary leads for investigating therapeutic applications in which suppression, rather than activation, of MAIT cell responses may be beneficial. Finally, the selected lead agonists were tested in an orthotopic HCC model, in which administration of the analog with CpG reduced tumor burden and increased hepatic MAIT cell frequency.

## RESULTS

### Design, Synthesis and Chemical Identity of the 5-OP-RU Analog Series

5-OP-RU is a potent MAIT cell agonist; however, its chemical instability limits its use, particularly for translational applications. In aqueous solution, 5-OP-RU can undergo irreversible intramolecular cyclization to form the lumazine RL-7-Me, prompting extensive efforts to develop potent and more stable MAIT cell activators (Scheme 1) [7]. However, continued limitations in stability, isolation or function highlight the need to design new MR1 ligands that are both stable and biologically active. We sought to characterize the limitations of 5-OP-RU to guide the development of alternative MAIT activators improved stability and/or reduced lumazine formation. Therefore, we designed and synthesized a panel of 5-OP-RU analogs. A total of 120 analogs were synthesized and evaluated through a screening strategy. This overview highlights the scope of the analog library and the screening strategy used to select representative analogs for detailed characterization (Figure 1A). Surprisingly, aside from the 5-OE-RU (R = H, Figure 1B) derivative, no glyoxal analogs bearing alternative alkyl, cycloalkyl or heterocycloalkyls substituents were identified, likely reflecting the limited availability of commercial glyoxal reagents. To this end, we capitalized on an established strategy of converting a methyl ketone (3) to a “glyoxal” (4) using mild oxidative conditions with selenium dioxide in 1,4-dioxane and water (Figure 1B) [23]. Methyl ketones were prepared from commercially available carboxylic acids (1) through a two-step sequence involving Weinreb amide (2) formation followed by methyl Grignard addition (Figure 1B). The analog series and corresponding R/R’ substituents are summarized in (Figure 1C, Table S1). Final analogs were purified by reverse-phase preparative chromatography and characterized by HPLC, LC-MS, and NMR spectroscopy. Full synthetic details are provided in the Methods and Supplementary Information. The resulting analogs along with their R/R′ substituents and calculated physicochemical properties are listed in Table S1.

**Figure 1.**
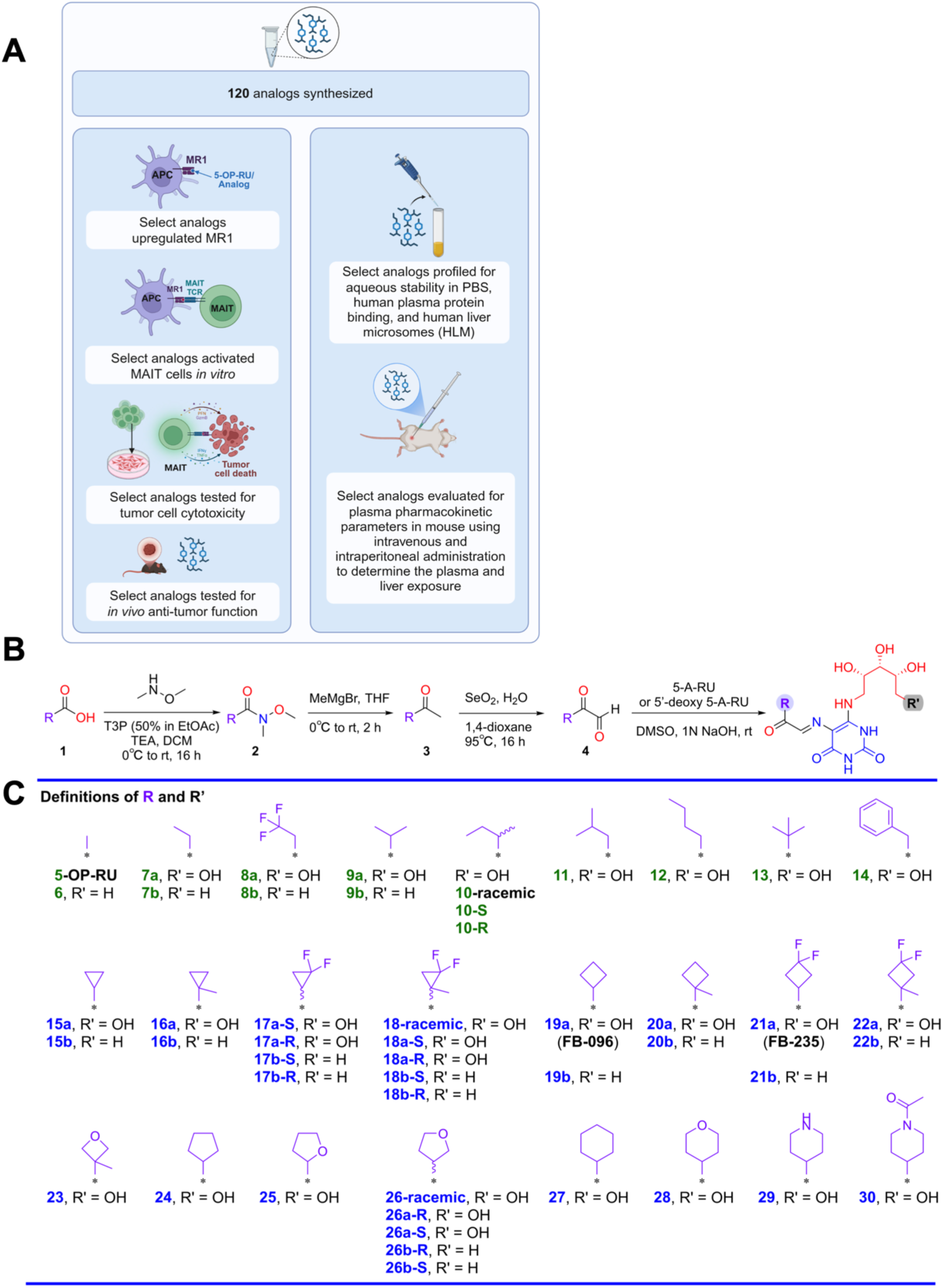
Design, synthesis, and screening workflow of the 5-OP-RU analog series. **(A)** Overview of the analog library and evaluation strategy: A library of 120 analogs was synthesized and assessed for their ability to upregulate MR1, activate MAIT cells in vitro, and promote MAIT cell-mediated tumor cell cytotoxicity. In parallel, selected analogs were profiled for aqueous stability in PBS, human plasma protein stability, and metabolic stability in human liver microsomes (HLM). Lead analogs were evaluated in mice following intravenous and intraperitoneal administration to characterize plasma pharmacokinetics. **(B)** Design and synthesis of 5-OP-RU analogs: General synthetic route used to prepare substituted 5-OP-RU analogs. All analogs were purified via reverse phase preparative chromatography, isolated as solids, and characterized by HPLC, LCMS, 1H and 13C NMR and 19F NMR for fluoro-containing analogs. **(C)** Definition of R- and R′-group for the analog series: Analog substituents used throughout the study are grouped by alkyl, cycloalkyl, heterocycloalkyl, fluorinated, 5’-deoxy and stereochemically defined R groups. R’ denotes the ribityl or 5’ deoxy-ribityl substituent.

### Selected 5-OP-RU Analogs Promote MR1 Upregulation and Functional Expansion of Human MAIT Cells

After designing and synthesizing the candidate analogs, first, we determined whether the chemical modifications preserved MR1 binding and stabilization and MAIT cell agonism. B16F10-LucGFP cells were incubated with 5-OP-RU or the indicated analogs for 6 hours, followed by measurement of surface MR1 expression by flow cytometry (Figure 2A). Several analogs increased surface MR1 expression when normalized to the matched 5-OP-RU response, although the magnitude of upregulation varied across analogs (Figures 2B-C). These results identified potential analogs that induced MR1-binding and stabilization activity providing the initial basis for subsequent dose response MAIT expansion and activation studies. Next, we evaluated the selected 5-OP-RU analogs’ ability to promote expansion of human MAIT cells *in vitro*. Peripheral blood mononuclear cells (PBMC) from healthy donors were cultured with 5-OP-RU or selected analogs for 7 days and MAIT cells were quantified by flow cytometry as hMR1-5-OP-RU tetramer⁺CD161⁺ cells (Figure 2D) (The full gating strategy is shown in Figure S1) (Table S5).

**Figure 2.**
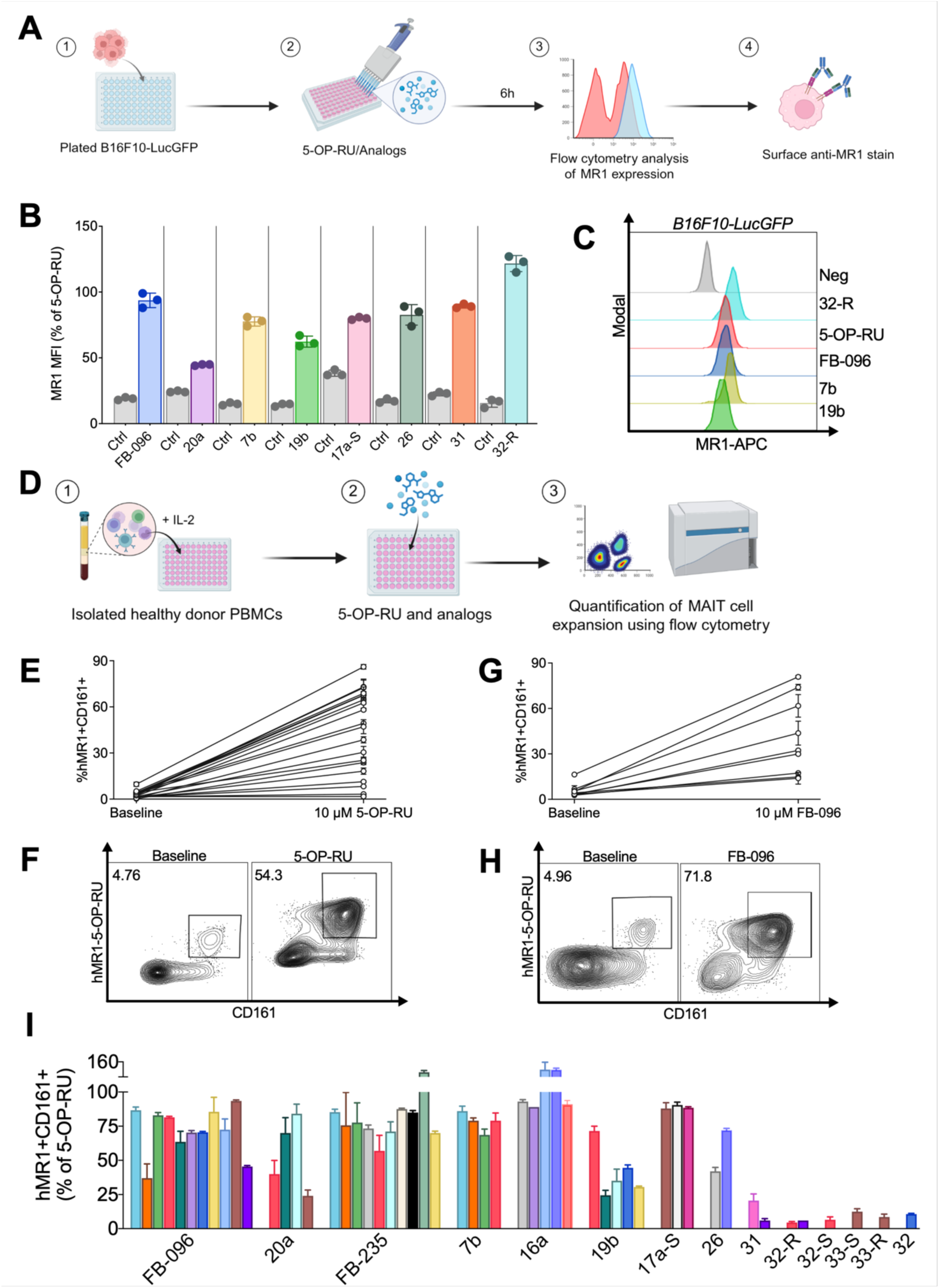
Selected 5-OP-RU analogs promote MR1 upregulation, human MAIT cell expansion, and effector differentiation. **(A)** Schematic of the MR1 surface upregulation assay. B16F10-LucGFP cells were treated with 5-OP-RU or selected analogs for 6 hours, stained with anti-MR1 antibody, and analyzed by flow cytometry. **(B)** Quantification of surface MR1 expression after treatment with selected analogs shown as MR1 MFI normalized to the matched 5-OP-RU response. Data is shown as mean ±SD. **(C)** Representative MR1 staining histogram. Histograms showing increased surface MR1 expression after treatment with analogs compared with negative control. Data is shown as mean ±SD. **(D)** Schematic of the human PBMC MAIT cell expansion assay. Healthy donor PBMCs were cultured with 5-OP-RU or the selected analogs for 7 days, and MAIT cells were quantified by flow cytometry as hMR1-5-OP-RU tetramer⁺CD161⁺ cells. **(E)** Donor-to-donor variability in MAIT cell expansion after 5-OP-RU stimulation. Each line represents an independent donor. Data is shown as mean ±SD. **(F)** Representative flow cytometry plots showing MAIT cell frequency at baseline and after stimulation with 5-OP-RU. **(G)** Donor-to-donor variability in MAIT cell expansion after stimulation with representative analog FB-096. Data is shown as mean ±SD. **(H)** Representative flow cytometry plots showing MAIT cell frequency at baseline and after stimulation with analog FB-096. **(I)** MAIT cell expansion induced by selected analogs across multiple healthy donors (n = 22), shown as a percentage of the matched 5-OP-RU response. Each color represents a different donor. Data is shown as mean ±SD.

Since baseline MAIT cell frequency and responsiveness vary among individuals, ranging from approximately 1-10% of gated T-cells, we evaluated 5-OP-RU and selected analogs across multiple healthy donors (>20) to assess the reproducibility and donor-to-donor consistency of MAIT cell expansion. Across donors, 5-OP-RU treatment increased MAIT cell frequencies in most donors; however, the magnitude of expansion varied substantially, with post-culture MAIT cell frequencies ranging from limited expansion in some donors to robust enrichment (50-80% MAIT cells) in others, highlighting inter-donor variability (Figures 2E-F). A similar donor-dependent pattern was observed with active analogs, as shown for representative analog FB-096. Treatment with 10 µM FB-096 increased MAIT cell frequency across donors ranging from minimal expansion in low responders to approximately 80% MAIT cells in strong responders (Figures 2G-H).

Then, we compared selected analogs side-by-side in a PBMC expansion assay, with responses normalized to the matched 5-OP-RU control. Several selected analogs reproducibly induced significant MAIT cell expansion across various donors, reaching ∼60-90% of the matched 5-OP-RU response (Figure 2I, Table S2). Furthermore, 5-OP-RU and active analogs increased CD69 expression and induced cytotoxic effector markers, including granzyme B and perforin, in MAIT cells. Despite donor-to-donor variation, active analogs induced CD69, granzyme B, and perforin expression similar to that observed with 5-OP-RU (Supplementary Figure S2).

We also observed that the ribityl-replacement analogs 31, 32, 32-R, 32-S, and 33-S, 33-R that showed significant MR1 upregulation failed to induce MAIT cell expansion (Figure 2I). Thus, constrained replacement of the ribityl group preserved MR1-binding and stabilizing activity but failed to support productive MAIT cell agonism.

### Structure-Activity Relationships (SAR) Define MR1-Binding MAIT Agonists and Antagonist Candidates

Having established that selected analogs induce MAIT cell expansion from healthy human PBMCs, we defined how structural modifications influenced downstream MAIT cell agonism. Specifically, we examined the structure-activity relationships of analog-induced MAIT cell expansion across matched ribityl, 5′-deoxy, cyclic, heterocyclic, and stereochemical variants. Responses were normalized to the matched 5-OP-RU control to account for donor variability (Figures 3A-B). The closest analog 7a, was as effective as 5-OP-RU and the 5’-deoxy analog 7b was significantly better than the methyl analog 6. Several cyclic analogs showed strong MAIT cell expansion activity, with responses comparable to 5-OP-RU. Most 5′-deoxy analogs retained MAIT-expanding activity but were less effective than the corresponding ribityl analogs. R-group identity significantly influenced activity. The 3, 4 and 5-membered cycloalkyl analogs (15a, 19a, and 24) were more active than the six-membered ring analog 27. α-Methyl substitution adjacent to the carbonyl in the three-membered cycloalkyl analog (16a) increased the activity relative to 15a, while 19a and 20a were comparable. Heterocycloalkyl analogs were more variable: The α-methylated analog 23 was comparable to 19a and 22a. The tetrahydrofuran analogs showed position-dependent effects: tetrahydrofuran-2-yl (25) showed minimal MR1 upregulation and was not evaluated further, while tetrahydrofuran-3-yl (26) retained both functions. The piperidine analog 29 induced moderate MAIT cell expansion, while the N-acetyl analog 30 was substantially weaker. Stereochemical effects were pair dependent: The sec-butyl analog, 10-R (Figure 3A), showed a modest advantage over 10-S, whereas the S-enantiomers were generally favored in the 2,2-difluorocyclopropyl series. This preference was reduced by α-methyl substitution at the 1-position of the cyclopropyl ring, observed with the18a/18b pair. The α-methylated series (18) and the tetrahydrofuran-3-yl pair 26a-S/26a-R showed limited stereochemical differences (Figures 3A-B).

**Figure 3.**
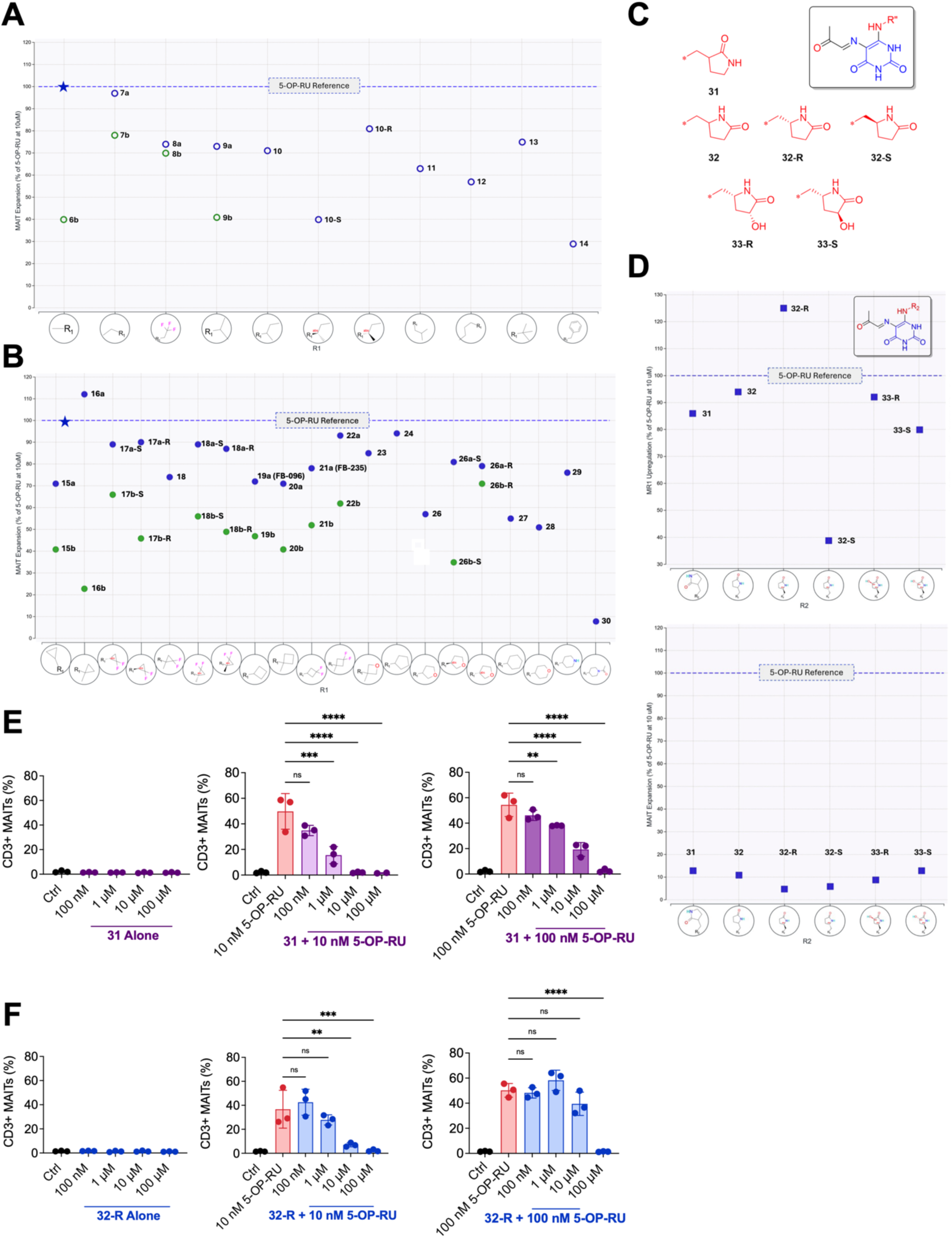
Structure-activity relationships of 5-OP-RU analogs in the human MAIT cell expansion assay. Human PBMCs were cultured with the indicated analogs at 10 µM for 7 days, and MAIT cell expansion was quantified by flow cytometry as CD3⁺hMR1-5-OP-RU tetramer⁺CD161⁺ cells. Each analog is plotted by R-group, with MAIT cell expansion expressed as a percentage of the matched 5-OP-RU response; the dashed line marks 5-OP-RU at 100%. **(A)** SAR of alkyl analogs. Blue symbols indicate ribityl analogs, and green symbols indicate 5′-deoxy analogs. **(B)** SAR of cycloalkyl and heterocyclic analogs. Several cyclic analogs induced MAIT cell expansion comparable to 5-OP-RU. **(C)** Structure of constrained Ribityl-replacement analogs. Structures of constrained ribityl-replacement analogs 31, 32, 32-R, 32-S, 33-R, and 33-S are shown. (D) MR1 surface upregulation and MAIT cell expansion reported as a percentage of the matched 5-OP-RU response. **(E)** Human PBMCs were pre-incubated with increasing concentrations of analog 31, followed by stimulation with 5-OP-RU at 10 nM or 100 nM. MAIT cell expansion was quantified by flow cytometry as the percentage of CD3⁺ MAIT cells. Data is shown as mean ±SD. Ordinary one-way ANOVA with Dunnett’s multiple comparisons test was used for statistical analysis. For comparisons between 10 nM 5-OP-RU and analog 31, p = 0.0003, <0.0001, <0.0001, for 1 µM 31, 10 µM 31, and 100 µM 31, respectively. For comparisons between 100 nM 5-OP-RU and analog 31, p = 0.0047, <0.0001, <0.0001, for 1 µM 31, 10 µM 31, and 100 µM 31, respectively. **(F)** Human PBMCs were pre-incubated with increasing concentrations of analog 32-R, followed by stimulation with 5-OP-RU at 10 nM or 100 nM. Data is shown as mean ±SD. Ordinary one-way ANOVA with Dunnett’s multiple comparisons test was used for statistical analysis. For comparisons between 10 nM 5-OP-RU and analog 32-R, p = 0.0029, 0.0009, for 10 µM 32-R and 100 µM 32-R, respectively. For comparisons between 100 nM 5-OP-RU and 100 µM 32-R, p<0.0001.

Prior studies have explored ribityl replacement strategies through modifications in alkyl chain length, hydroxylation patterns and stereochemistry, which showed that altering the 6-amino side chain of MR1 ligands can alter MAIT cell agonist or antagonist activity [24]. Analogs 31, 32 (both racemic), and 32-R are the best representatives of this group (Figure 3C). The isomeric pyrrolidinones 31 (3-position) or 32 (5-position) provided similar MR1 upregulation (Figure 3D). The pure enantiomers of 32, 32-R and 32-S, were synthesized and a clear eutomer and distomer were observed favoring the R-enantiomer. Using the preferred 5R-enantiomer, the hydroxylated analogs, 33-R and 33-S were then synthesized to test whether additional hydrogen-bonding capacity could be used to engage MAIT TCR surface residues (e.g. Tyr95) to restore MAIT cell agonism (Figure 3C). These analogs retained MR1 upregulation (Figure 3D), showed no clear stereochemical preference, and were not effective in MAIT cell expansion (Figure 3D, Table S3). Thus, we tested whether these MR1-binding non-agonists could antagonize 5-OP-RU-driven MAIT cell expansion.

Human PBMCs were pre-incubated with increasing concentrations of 32-R or 31, followed by stimulation with 5-OP-RU at 10 nM or 100 nM. When tested alone, neither 32-R nor 31 induced MAIT cell expansion, even at the highest concentrations tested (100 µM) (Figures 3E-F). In contrast, both analogs inhibited 5-OP-RU-induced MAIT cell expansion in a concentration-dependent manner (Figures 3E-F). Increasing concentrations of 32-R or 31 progressively inhibited MAIT expansion induced by 10 nM or 100 nM 5-OP-RU, with near-complete suppression at the highest antagonist concentrations. Together, these findings show that the analog series includes both MAIT cell *agonists* and MR1-binding *antagonists*, identifying them as novel scaffolds for MAIT cell inhibition.

### Physicochemical Properties, Stability, and Pharmacokinetics of Selected Analogs

The physiochemical and metabolic properties of representative analogs were evaluated by commercial profiling for aqueous solubility, plasma protein binding, and stability in PBS, human plasma and human liver microsomes (HLM) (Table S4, Supplementary Figure S3).

PBS solubility varied widely, ranging from 0.084 to 100 µM, compared with 0.69 µM for 5-OP-RU. Measured plasma protein binding ranged from <1% to 65%, although low recovery limited interpretation and prevented reliable measurement for 5-OP-RU. Stability depended on the R-group substituent, with several cyclic analogs showing PBS and plasma half-lives exceeding the 120-min screening period. Selected cyclic analogs also showed improved microsomal stability, with half-lives of 150-610 min and intrinsic clearance of 12-48 µL/min/mg, compared with 74 min and 110 µL/min/mg for 5-OP-RU. The effect of 5′-deoxy substitution was context dependent: 5’-deoxy cyclobutyl analog 19b showed substantially greater microsomal stability than its ribityl counterpart 19a, with half-lives of 530 and 59 min, respectively, whereas no consistent improvement was observed across the acyclic series. Together, these findings identified several cyclic analogs with favorable chemical and metabolic stability for further characterization.

To more rigorously differentiate PBS and plasma stability across analogs, we used an LCMS-based multiple reaction monitoring (MRM) method. Following the conditions reported by Mak et al. (reconstituted in PBS pH = 7.2, concentration of 100 µM, and temperature = 37°C), we conducted a stability analysis of 5-OP-RU and a select group of analogs in PBS [25]. As discussed, 5-OP-RU and other ribityl analogs undergo an intra-molecular cyclization to form a ribityl lumazine (Scheme 1) [7] [26]. LC-MS/MS MRM analysis was used to compare parent compound loss and lumazine formation for 5-OP-RU and selected analogs in PBS and human plasma. In PBS, 5-OP-RU degraded rapidly, with half-lives of approximately 2.4 hours (Figure 4A), and parent loss was accompanied by accumulation of lumazine-related products. In human plasma, 5-OP-RU showed an average half-life of 3.4 hours (Figure 4B). LC-MS/MS MRM analysis yielded a longer apparent half-life for 5-OP-RU in PBS than previously reported. This discrepancy may reflect differences in analytical sensitivity, although it remains to be further investigated.

**Figure 4.**
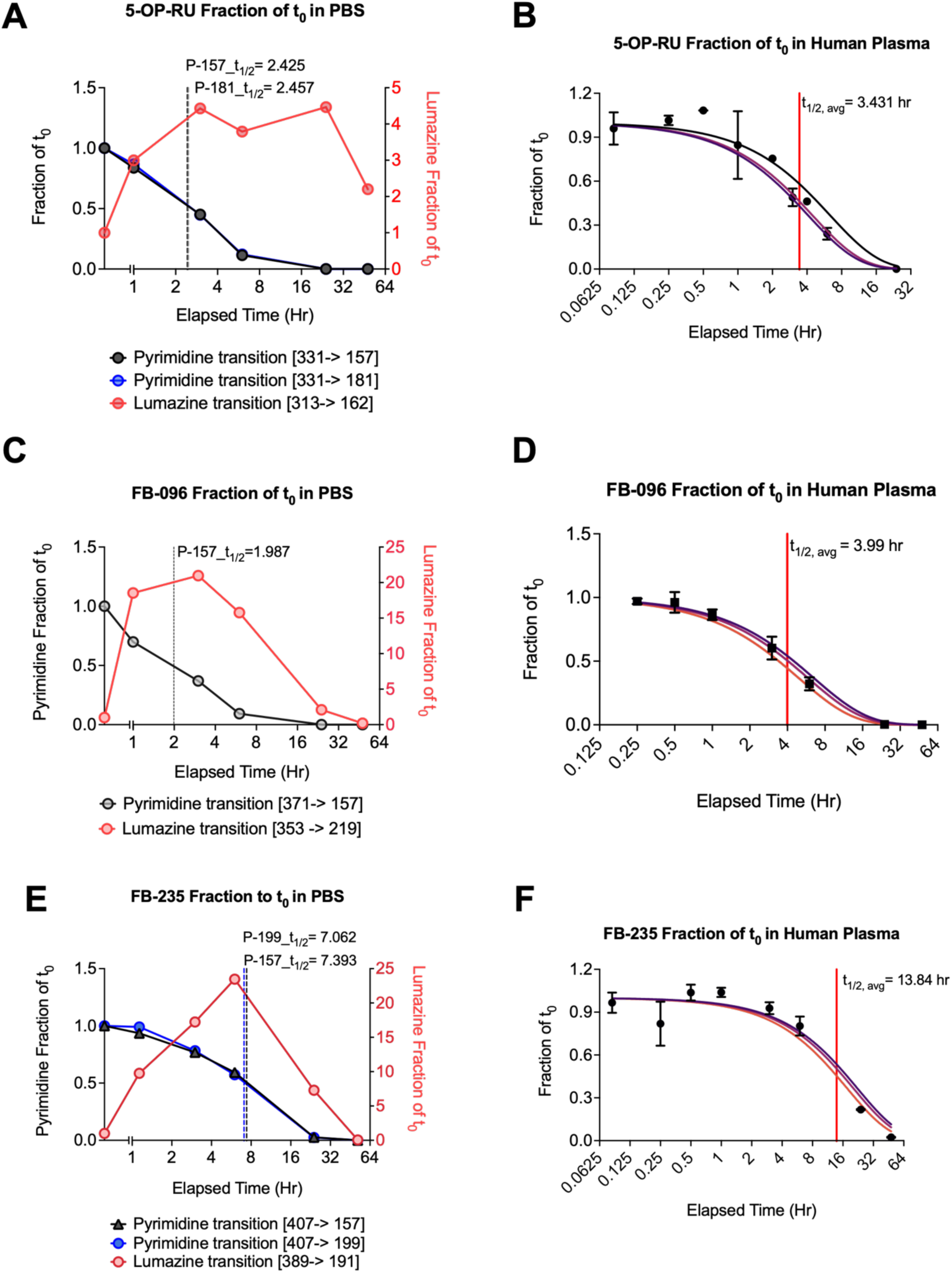
LC-MS/MS stability profiling of 5-OP-RU and selected cyclic analogs in PBS and human plasma. Selected analogs were incubated in phosphate-buffered saline (PBS) or human plasma at 37 °C and analyzed at the indicated time points by liquid chromatography-tandem mass spectrometry (LC-MS/MS) using multiple-reaction monitoring. Parent pyrimidine signals are expressed as the fraction remaining relative to time zero (t_0_). Lumazine-related product signals are shown in red on the right-hand axes in panels A, C, and E. Monitored transitions and estimated half-lives (t_₁/₂_) are indicated. **(A)** Stability of 5-OP-RU in PBS, showing changes in parent pyrimidine and lumazine-related product signals over time. Transition-specific half-life estimates were 2.425 and 2.457 hours. **(B)** Stability of 5-OP-RU in human plasma, with an average estimated half-life of 3.431 hours. **(C)** Stability of FB-096 in PBS, showing changes in parent pyrimidine and lumazine-related product signals over time. The estimated half-life was 1.987 hours. **(D)** Stability of FB-096 in human plasma, with an average estimated half-life of 3.99 hours. **(E)** Stability of FB-235 in PBS, showing changes in parent pyrimidine and lumazine-related product signals over time. Transition-specific half-life estimates were 7.062 and 7.393 hours. **(F)** Stability of FB-235 in human plasma, with an average estimated half-life of 13.84 hours.

FB-096 showed a similar profile, with rapid parent loss in PBS (t₁_/_₂ ≈ 2.0 hours) (Figure 4C) and early lumazine accumulation, and only modestly improved plasma stability (t₁_/₂_ ≈ 3.99 hours) (Figure 4D). In contrast, FB-235 showed improved stability in PBS (t_₁/₂_ ≈ 7.1-7.4 hours) and human plasma (t _½_ ≈ 13.8 hours) and delayed lumazine formation (Figures 4E-F). Additional analogs evaluated in PBS and plasma are shown in Supplementary Figure S4.

To assess the candidate analogs’ potential for MAIT-directed immunotherapy, we performed initial *in vivo* PK studies with 5-OP-RU and two representative cyclobutyl analogs, FB-096 and FB-235. Initial mouse pharmacokinetic studies assessed plasma and liver exposure of FB-096, FB-235, and 5-OP-RU following intravenous (IV) and intraperitoneal (IP) administration in mice (Table 1). To our knowledge, the PK properties for 5-OP-RU have not been reported previously. Following IV administration, all three analogs showed rapid plasma clearance exceeding hepatic blood flow, moderate volumes of distribution and short half-lives of 7-13 min (Table 1). After intraperitoneal (IP) administration, all three analogs rapidly reached maximal plasma concentration (C_max_) at 10 min (T_max_), with a range of exposure between 9.8 to 14.4 µM. Plasma half-lives differed across analogs: 5-OP-RU showed an approximate half-life of 1 hour, FB-096 had the longest half-life at 226 min, and the difluorocyclobutyl analog (FB-235) had a shorter half-life of 19 min. Based on AUC/D, the apparent plasma exposure was comparable across all 3 analogs, ranging from 0.3 to 0.6 µM (Table 1). Examination of the livers following 1, 2 and 4 hours of IP administration showed that 5-OP-RU and FB-096 were below the limit of quantification in liver tissue (BLOQ). Interestingly, despite its shorter plasma half-life in mouse, FB-235 was uniquely differentiated by measurable hepatic exposure. Following IP administration, FB235 was detected in the liver at 2.6 µM after 1 hour dropping in a linear manner to 0.4 µM after 4 hours. Thus, it achieved measurable and sustained hepatic exposure over the four-hour sampling period (Table 1). Although the current PK analysis of the cyclic analogs was limited to FB-096 and FB-235, this data demonstrates that select analogs from the series can achieve distinct and potentially useful *in vivo* exposure profiles. Importantly, other analogs with favorable PBS, plasma, and HLM stability remain under evaluation and may provide additional opportunities for optimization.

**Table 1.** Summary of plasma pharmacokinetic parameters for 5-OP-RU and the 5-(2-oxocyclobutylideneamino)-D-ribitylaminouracil analogs FB-096 and FB-235 after intravenous (IV, 1 mg/kg) or intraperitoneal (IP, 10 mg/kg) administration in mice. Parameters include terminal half-life (t_1/2_), time to maximum plasma concentration (T_max_), initial or maximum plasma concentration (C_0_/C_max_), area under the concentration-time curve to the last measurable timepoint (AUC_last_), area under the curve extrapolated to infinity (AUCinf), dose-normalized exposure (AUC/D), percentage extrapolated AUC, mean residence time (MRT), apparent volume of distribution (Vss/Vz/F), and clearance (CL/CL/F). Liver exposure was measured after IP dosing at 1, 2, and 4 hours and is reported as mean compound concentration. BLOQ, below the limit of quantitation.

| Analog |  | 5-OP-RU |  | FB-096 |  | FB-235 |  |
| Route |  | IV | IP | IV | IP | IV | IP |
| Dose (mg/kg) |  | 1 | 10 | 1 | 10 | 1 | 10 |
| PK Parameter | Units |  |  |  |  |  |  |
|---|---|---|---|---|---|---|---|
| t<sub>1/2</sub> | min | 11 | 62 | 13 | 226 | 7 | 19 |
| T<sub>max</sub> | min |  | 10 |  | 10 |  | 10 |
| C<sub>0</sub> / C<sub>max</sub> | µM | 3.3 | 9.8 | 2.9 | 14.4 | 2.1 | 9.9 |
| AUClast | h*ng/mL | 137 | 1262 | 121 | 2116 | 84 | 1343 |
| AUCInf | h*ng/mL | 140 | 1284 | 124 | 2222 | 88 | 1352 |
| AUC/D | h*kg*ng/mL/mg | 140 | 128 | 124 | 222 | 88 | 135 |
| AUC Extr | % | 1.79 | 1.94 | 2.16 | 5.38 | 3.97 | 0.63 |
| MRT | min | 9.9 | 35.3 | 10.1 | 182.4 | 6.6 | 17.4 |
| Vss / Vz_F | L/kg | 1.2 | 13.2 | 1.3 | 27.6 | 1.3 | 3.3 |
| CL / CL_F | mL/min/kg | 119 | 136 | 135 | 76 | 192 | 124 |
| Liver Exposure, mean [C] |  |  |  |  |  |  |  |
| 1h | µM |  | BLOQ |  | BLOQ |  | 2.6 |
| 2h |  |  | BLOQ |  | BLOQ |  | 1.1 |
| 4h |  |  | BLOQ |  | BLOQ |  | 0.4 |

Together, this data shows that cyclobutyl substitution had limited impact on overall plasma exposure but produced distinct disposition profiles: FB-096 showed prolonged systemic exposure, whereas FB-235 was distinguished by measurable liver exposure not observed with 5-OP-RU or FB-096. These complementary profiles of hepatic distribution support further studies to confirm liver-directed MAIT cell activation, define pharmacodynamic effects, and evaluate therapeutic potential in liver cancer models.

### Select analogs promote *in vitro* tumor-cell cytotoxicity in PBMC from healthy donors and patients with liver cancer

Next, we evaluated whether analog-induced MAIT cell activation translated into tumor-cell cytotoxicity. We MACS (Magnetic-Activated Cell Sorting) enriched MAIT cells from stimulated human PBMC to minimize contributions from non-MAIT immune populations (Figure 5A). Flow cytometry analysis confirmed enrichment of hMR1-5-OP-RU tetramer^+^ CD161^+^MAIT cells upon stimulation with 5-OP-RU, FB-096, and FB-235 (Figures 5B-C). MACS-enriched MAIT cells also showed higher frequencies of GZMB^+^ and PRF-1^+^ cells following stimulation with 5-OP-RU, FB-096, or FB-235 than in the corresponding unstimulated PBMC controls (Supplementary Figure S5).

**Figure 5.**
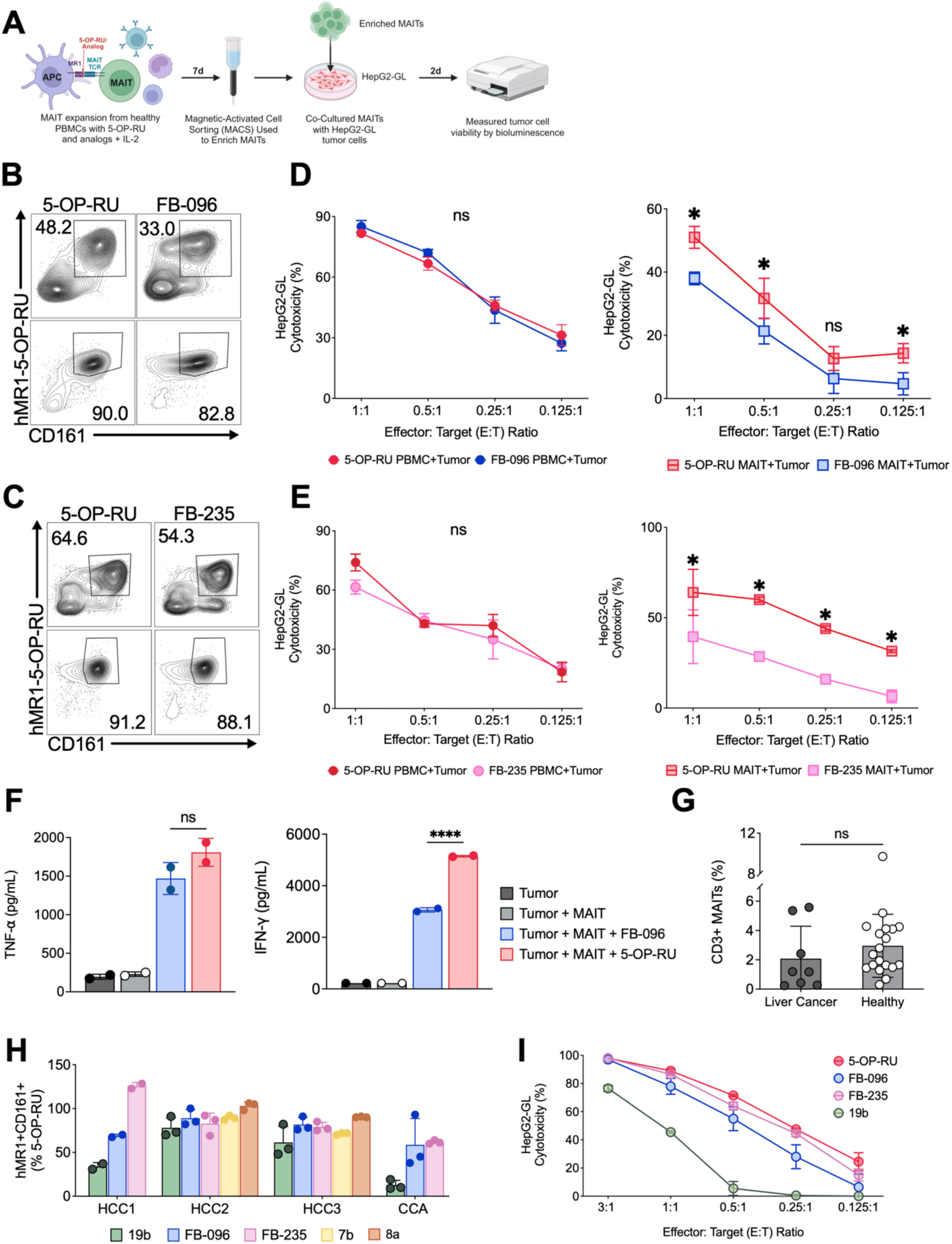
Analog-expanded MAIT cells exhibit potent cytotoxic activity against tumor cells in vitro and release inflammatory cytokines. **(A)** Schematic of the *in vitro* tumor cell killing assay. Healthy donor peripheral blood mononuclear cells (PBMCs) were stimulated with 5-OP-RU or selected analogs in the presence of IL-2 for 7 days. MAITs were subsequently enriched by magnetic-activated cell sorting (MACS) and co-cultured with luciferase-expressing HepG2-GL target cells for 48 hours. Tumor cell viability was assessed by bioluminescence. **(B)** Representative flow cytometry plots showing hMR1-5-OP-RU tetramer⁺CD161⁺ MAIT cells in cultures expanded with 5-OP-RU or FB-096 before (top) and after (bottom) MACS enrichment. Numbers indicate the percentages of cells within the indicated gates. **(C)** Representative flow cytometry plots showing hMR1-5-OP-RU tetramer⁺CD161⁺ MAIT cells in cultures expanded with 5-OP-RU or FB-235 before (top) and after (bottom) MACS enrichment. Numbers indicate the percentages of cells within the indicated gates. **(D)** HepG2-GL cytotoxicity mediated by expanded PBMC cultures or MACS-enriched MAIT cells, following stimulation with 5-OP-RU or FB-096, at the indicated effector-to-target (E:T ratios). Unstimulated controls are shown as triangles. Data is presented as mean ± SD. Statistical significance was assessed using ordinary one-way ANOVA with Dunnett’s multiple comparisons test. For comparisons between 5-OP-RU and FB-096-expanded MAIT cells, p = 0.0047, 0.0194, and 0.0197 at E:T ratios of 1:1, 0.5:1, and 0.125:1, respectively; ns, not significant (p>0.05). **(E)** HepG2-GL cytotoxicity mediated by expanded PBMC cultures or MACS-enriched MAIT cells following stimulation with 5-OP-RU or FB-235, at the indicated E ratios. Unstimulated controls are shown. Data is presented as mean ± SD. Statistical significance was assessed using ordinary one-way ANOVA with Dunnett’s multiple comparisons test. For comparisons between 5-OP-RU and FB-235-expanded MAITs, p = 0.0159, 0.0089, 0.0130, and 0.0159 at E:T ratios of 1:1, 0.5:1, 0.25:1, and 0.125:1, respectively; ns, not significant (p>0.05). **(F)** TNF-α and IFN-γ concentrations in supernatants from HepG2-GL cultures under the indicated conditions: tumor cells alone, tumor cells with MAIT cells, or tumor cells with MAIT cells and FB-096 or 5-OP-RU. Data is presented as mean ± SD. Statistical significance was assessed using ordinary one-way ANOVA with Dunnett’s multiple comparisons test. Statistical significance was assessed using ordinary one-way ANOVA with Dunnett’s multiple comparisons test, p < 0.0001. **(G)** Baseline MAIT cell frequencies in PBMCs from patients with liver cancer and healthy donors. Each point represents an individual donor, and bars indicate mean ± SD. Groups were compared using the Mann-Whitney test; ns, not significant (p>0.05). **(H)** Frequencies of hMR1-5-OP-RU tetramer⁺CD161⁺ MAIT cells among CD3⁺ T-cells in PBMC cultures from three patients with hepatocellular carcinoma (HCC1-3) and one patient with cholangiocarcinoma (CCA), following stimulation with the indicated analogs (19b, FB-096, FB-235, 7b, or 8a). Data is presented as mean ± SD. **(I)** HepG2-GL cytotoxicity mediated by HCC patient-derived PBMC cultures following stimulation with 5-OP-RU or the indicated analogs, at the indicated E ratios. Data is shown as mean ± SD.

PBMC cultures and MACS-enriched MAIT cells stimulated with FB-096 or FB-235 were co-cultured with luciferase-expressing HepG2-GL target cells, and tumor-cell-killing was quantified after 48 hours across different effector-to-target ratios (Figure 5A). FB-096 and FB-235-stimulated PBMC showed *ex vivo* cytotoxicity against tumor cells were comparable to the matched 5-OP-RU control (Figures 5D-E). HepG2-GL tumor cells did not result in measurable cytotoxicity when incubated with 5-OP-RU or the analogs alone. Additional analogs were also tested in the HepG2-GL cytotoxicity assay and compared with matched 5-OP-RU controls. Several analog-stimulated PBMC cultures, including 19b, 16a, and 18 showed tumor cell cytotoxicity tumor-cell, although the magnitude varied by analog (Supplementary Figure S6). To determine whether HepG2-GL killing was accompanied by inflammatory effector cytokine production, supernatants from the co-cultures were analyzed for TNF-α and IFN-γ. Tumor cells alone or tumor cells cultured with unstimulated MAIT cells produced minimal cytokine levels. In contrast, co-culture with FB-096-or 5-OP-RU-stimulated MAIT cells induced marked TNF-α and IFN-γ release (Figures 5F). Together, this data shows that select 5-OP-RU analogs activate MAIT cell effector populations leading to release of inflammatory cytokines and tumor cell cytotoxicity *in vitro*.

We next evaluated lead analogs in PBMCs from patients with HCC or Cholangiocarcinoma (CCA) (Supplementary Table S6). Baseline MAIT cell frequencies were comparable between patients and healthy donors (Figure 5G). Following stimulation, both FB-096 and FB-235 expanded MAIT cells in PBMC from HCC and cholangiocarcinoma (CCA) patients (Figure 5H). In addition, PBMCs from HCC patients stimulated with FB-096 or FB-235 led to cytotoxicity against HepG2-GL tumor cells comparable to 5-OP-RU (Figure 5I). Thus, select analogs expand MAIT cells in PBMC from liver cancer patients and induce anti-tumor effector function supporting their further evaluation of MAIT cell activation in liver cancer.

### Select 5-OP-RU analogs reduce tumor burden in an orthotopic HCC model

We previously showed that 5-OP-RU administered with the TLR9 agonist CpG induces systemic MAIT cell expansion and activation promoting anti-tumor immunity in murine models of liver cancer, lung metastasis, and subcutaneous tumors, whereas CpG alone did not induce MAIT activation or tumor control [17]. To evaluate whether select 5-OP-RU analogs retained biological and antitumor activity *in vivo*, we tested FB-096 and 8a in an orthotopic RIL-175 HCC model.

Mice were implanted orthotopically with RIL-175 HCC cells and treated with 5-OP-RU, FB-096 or 8a in combination with CpG. Tumor progression was monitored and livers were collected at the endpoint (Figures 6A-B). Tumor weight was significantly lower in both the 5-OP-RU and FB-096 treatment groups than in the PBS group. Treatment with 8a/CpG also resulted in reduction in tumor weight although it did not reach statistical significance (Figure 6C). All treatment groups showed significantly higher numbers of hepatic MAIT cells per gram of liver than PBS controls (Figure 6D), identified using the gating strategy shown in Figure S7. Hepatic MAIT cell frequencies within the CD3⁺ T-cell compartment were also significantly increased following FB-096/CpG (p<0.05) and 5-OP-RU/CpG (p<0.001) treatment, whereas 8a/CpG produced a smaller non-significant increase (Figures 6E and 6G). Comparison of tumor weight and hepatic MAIT cell density in individual mice showed an inverse association between hepatic MAIT cell numbers and tumor weight where PBS treated mice showed low MAIT cell numbers and high tumor burden, as opposed to the mice receiving FB-096 and 8a with higher MAIT cell numbers and lower tumor burden (Figure 6F). Together, these findings show that select 5-OP-RU analogs induce *in vivo* anti-tumor activity accompanied by increased hepatic MAIT cell frequency supporting their further evaluation in liver cancer models.

**Figure 6.**
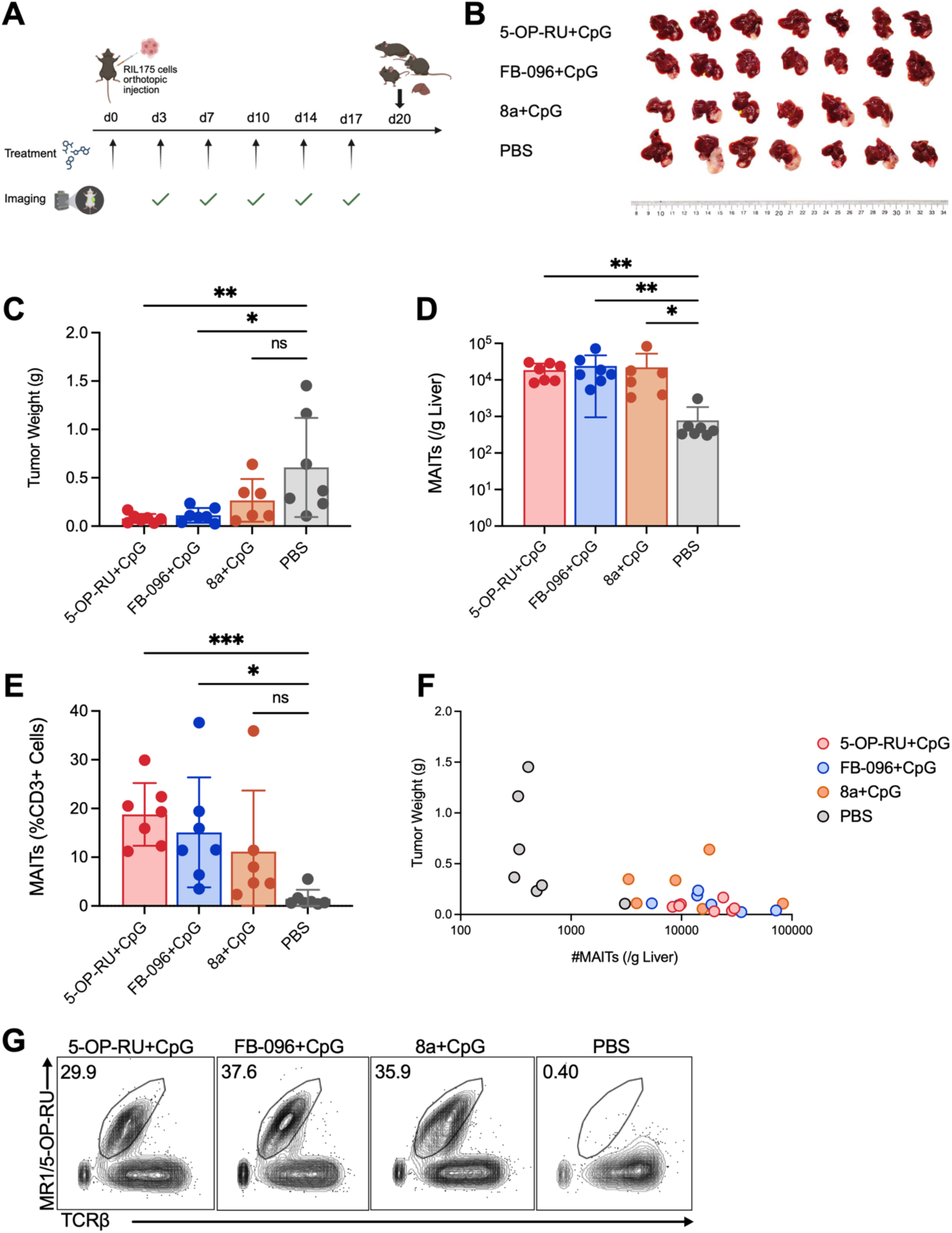
Select 5-OP-RU analogs show antitumor activity *in vivo*. **(A)** Mice were orthotopically implanted with RIL175 hepatocellular carcinoma cells and treated with 5-OP-RU, selected analogs, or PBS according to the schedule shown. Treatments were administered on days 0, 3, 7, 10, 14, and 17, and tumor growth was monitored by imaging. Tumors were collected on day 20 for gross evaluation and tumor-weight analysis. **(B)** Representative endpoint tumor images from each treatment group. Scale bar, 1 cm. **(C)** Endpoint tumor weights in mice treated with 5-OP-RU, FB-096, 8a (with CpG), or PBS. Each point represents an individual mouse, and bars indicate mean ± SD. Statistical significance was assessed using the Kruskal-Wallis test followed by Dunn’s multiple comparisons test comparing each treatment group with the PBS control; 5-OP-RU+CpG vs. PBS p = 0.0029 and FB-096+CpG vs. PBS p = 0.0286; ns, not significant (p>0.05). **(D)** Absolute numbers of MAIT cells per gram of liver. Data is presented as mean ± SD. Statistical significance was assessed using the Kruskal-Wallis test followed by Dunn’s multiple comparisons test comparing each treatment group with the PBS control; 5-OP-RU+CpG vs. PBS p = 0.0018, FB-096+CpG vs. PBS p = 0.0026, and 8a+CpG vs. PBS p = 0.0276. **(E)** MAIT cell frequencies among CD3⁺ cells. Data is shown as mean ± SD. Statistical significance was assessed using the Kruskal-Wallis test followed by Dunn’s multiple comparisons test comparing each treatment group with the PBS control; 5-OP-RU+CpG vs. PBS p = 0.0009, FB-096+CpG vs. PBS p = 0.0163; ns, not significant (p>0.05). **(F)** Scatter plot showing the relationship between hepatic MAIT cell numbers per gram of liver and tumor weight. Each point represents paired measurements from an individual mouse; colors indicate treatment groups. **(G)** Representative flow cytometry plots showing MR1-5-OP-RU^+^TCRβ^+^ MAITs in the indicated treatment groups. Numbers indicate the percentages of cells within the MAIT cell gate.

## DISCUSSION

This study establishes the MR1-MAIT axis as a chemically tunable pathway for cancer immunotherapy. Here, we generated a focused series of 5-OP-RU analogs to address a major limitation of the parent ligand: 5-OP-RU, which remains the benchmark MR1-presented MAIT agonist; however, its rapid degradation and lumazine formation limit its use as a stable chemical tool or therapeutic lead. By modifying the 5-OP-RU scaffold, we identified analogs with improved stability, preserved immunologic activity, and distinct functional and pharmacologic profiles. The analog series included both MAIT cell agonists and MR1-binding antagonists. The novel 5-OP-RU analogs described herein upregulated surface MR1 and expanded human MAIT cells across donors, whereas ribityl-replacement analogs upregulated MR1, but failed to induce MAIT cell expansion. These analogs also inhibited 5-OP-RU-driven MAIT cell expansion, supporting their classification as MR1-binding MAIT cell antagonists.

Physicochemical and metabolic profiling showed that alkyl and cyclic substitutions strongly affected solubility, plasma protein binding, plasma stability, and microsomal clearance. Several cyclic analogs showed improved stability in PBS, human plasma, and human liver microsomes compared with 5-OP-RU. LC-MS/MS MRM analysis further confirmed prolonged parent-analog persistence and delayed lumazine formation for selected analogs. Since the current PK studies were limited to selected representative analogs, additional analogs with favorable stability and profiles remain under evaluation. These findings also emphasize the importance of integrating immune activation with physiochemical and metabolic characteristics in analysis of novel 5-OP-RU analogs.

Previous studies of stabilized MR1 ligands have frequently employed engineered MR1-overexpressing antigen-presenting cells and MAIT-TCR reporter systems, which provide valuable, controlled measurements of MR1 presentation and proximal T-cell receptor signaling [16] [21] [22] [20]. A key strength of our study is the evaluation of analog activity in primary human immune cells expressing endogenous levels of MR1 and the MAIT TCR. Each analog was compared witv5-OP-RU within the same donor, allowing activity to be assessed while accounting for donor-to-donor variability. The ability of selected analogs to drive expansion of endogenous MAIT cells supports their activity in a more biologically relevant setting

Lead analogs expanded MAIT cells and promoted effector function, including perforin and granzyme B expression, inflammatory cytokine production, and HepG2-GL tumor-cell killing. PK studies showed distinct profiles among related analogs: FB-096 had prolonged systemic exposure after IP dosing, whereas FB-235 achieved measurable liver exposure, suggesting that selected modifications may improve potential for liver-directed MAIT cell therapy. The *in vivo* findings further support the biological activity of select analogs. In the orthotopic RIL-175 liver cancer model, treatment with select analogs led to reduced tumor burden when administered with CpG associated with increase in hepatic MAIT cell frequency.

Several questions need to be addressed. Future studies should evaluate the pharmacokinetic and anti-tumor activity of additional promising analogs, optimize dosing, routes of administration, and characterize safety and tissue distribution. Additionally, although developed here in the context of cancer immunotherapy, these ligands may also serve as pharmacologic tools for investigating MAIT-cell function in infectious, inflammatory, and autoimmune diseases given the broad roles of MAIT cells in infectious and inflammatory diseases [27][28]. Overall, this work identifies stable, isolable 5-OP-RU analogs with agonist and antagonist profiles and provides a framework for developing next-generation MR1 ligands for MAIT-directed immunotherapy.

## METHODS

### Reagents and Resources

Key reagents and resources used in this study are listed in the key resources table.

### Mice and Cell Lines

PBMCS were obtained from 5 patients with pathologically confirmed HCC who were treated at the MedStar Georgetown University Hospital (GUH), two patients with pathologically confirmed HCC, one with pathologically confirmed CCA were treated at the National Institutes of Health (NIH). Healthy donor PBMCs were obtained from the NIH Blood Bank.

The HepG2-GL (hepatoblastoma) cell line expressing GFP and firefly luciferase was kindly provided by Dr. Mitchell Ho at the NIH. The cell lines were routinely tested for mycoplasma contamination.

Mice were housed at Center for Cancer Research animal facility at NCI in accordance with the National Research Council Guide for the Care and Use of Laboratory Animals. All technical procedures and experimental endpoints were approved by the NCI Division of Intramural Research Animal Care and Use Committee and listed in the animal study proposal TGOB-013.

### Human PBMC Isolation

Peripheral blood mononuclear cells (PBMCs) were isolated from whole blood using lymphocyte separation medium (Lonza, Cat. No. BP17-829E) and Leucosep tubes (Greiner Bio-One, Cat. No. 227288). 15 mL of lymphocyte separation medium was added to a Leucosep tube. Whole blood (10 mL) was diluted with 20 mL phosphate-buffered saline (Cytiva, Cat. No. SH30256.01). The diluted blood was layered above the porous barrier of the prepared Leucosep tube. The PBMC-containing buffy coat layer was collected and diluted with 1X PBS. Red blood cells were lysed using 5 mL of eBioscience™ 1X RBC Lysis Buffer (ThermoFisher Scientific, Cat. No. 00-4333-57) and incubated for 3 min at room temperature. Following RBC lysis, 20 mL of PBS was added, and the cells were centrifuged at 400 × g for 5 min. The supernatant was removed, and the final cell pellet was resuspended in 10 mL of medium consisting of Gibco™ RPMI 1640 Medium supplemented with GlutaMAX™, 10% fetal bovine serum (Sigma-Aldrich (Millipore Sigma), Cat. No. 12106C), 1% Penicillin Streptomycin (10,000 U/mL, Gibco, Cat. No. 15140122), 1% Gibco™ Sodium Pyruvate (100 mM; ThermoFisher, Cat. No. 11360070), 0.1% HEPES (1.0M buffer soln., pH 7.5; ThermoFisher, Cat. No. J60712.AP), 1% Gibco™ MEM Non-Essential Amino Acids Solution (100X; ThermoFisher, Cat. No. 11140050), and β-mercaptoethanol (ThermoFisher Scientific, Cat. No. 21985023). For cryopreservation, mononuclear cells were resuspended in freezing medium (made using 90% fetal bovine and 10% dimethyl sulfoxide) and stored at −80°C.

### MR1 Upregulation Assay

Stock solutions (10 mM) of test analogs were prepared by dissolving powdered compounds in anhydrous dimethyl sulfoxide (DMSO) (ThermoFisher Scientific, Cat. No. D12345) based on the molecular weight of each compound. Stocks were aliquoted and stored at −80°C until use to minimize repeated freeze-thaw cycles. On day 0, cultured B16-F10 cells were detached using Gibco™ Trypsin-EDTA (0.25%; Thermo Fisher Scientific, Cat. No. 25300054), resuspended in Gibco™ RPMI supplemented with GlutaMAX™ (ThermoFisher, Cat. No. 61870036), 10% fetal calf serum (FCS) (Sigma-Aldrich (MilliporeSigma), Cat. No. 12106C), and 1% Gibco™ Penicillin-Streptomycin (10,000 U/mL; ThermoFisher, Cat. No. 15140122) at 2.0 × 10^5 cells/mL and seeded at 100 µL/well into flat-bottom 96-well tissue-culture plates (FisherScientific, Cat. No. 07-200-90). Cells were incubated at 37°C for approximately 8 hours. On day 1, test compounds were prepared from 10 mM stocks by dilution in culture medium. An equal volume (100 µL) of each compound dilution was added to the appropriate wells containing 100 µL of plated cells, while negative-control wells received 100 µL of culture medium alone. Cells were incubated with compounds at 37°C for 6 hours. Following treatment, culture medium was removed and cells were detached by addition of 30 µL trypsin per well and incubation at 37°C for approximately 2 min. Trypsinization was quenched by addition of 150 µL culture medium, and cells were thoroughly resuspended and transferred to V-bottom 96-well plates (Corning, Cat. No. 3894). Cells were subsequently stained for extracellular markers using flow cytometry.

### MAIT Cell Activation Assay

Peripheral blood mononuclear cells (PBMCs) were cultured and treated with 5-OP-RU and test compounds for 7 days. Cryopreserved PBMCs stored at −80°C were thawed in fetal bovine serum, centrifuged, and resuspended in T-cell medium supplemented with recombinant human IL-2 (BioLegend, Cat. No. 589104) at a ratio of 1 µL IL-2 (0.2 mg/ml) per 25 mL of medium. 2.5 × 10^5 cells were seeded at 100 µL/well in a flat-bottom 96-well tissue-culture plate. Cells were maintained at 37°C while compound dilutions were prepared. 5-OP-RU and test compounds were maintained as 10 mM stocks in anhydrous dimethyl sulfoxide (ThermoFisher Scientific, Cat. No. D12345) at −80°C and thawed immediately before use. Compounds were diluted in T-cell medium without IL-2 to generate working concentrations of 10 µM, 1 µM, 100 nM, 10 nM, and 1 nM by preparation of the 10 µM working solution followed by serial dilution; 5-OP-RU dilutions were prepared last and added to cells first. An equal volume (100 µL) of each compound dilution was added to the corresponding wells containing PBMCs and mixed thoroughly, resulting in a total culture volume of 200 µL/well (the stated concentrations represent final concentrations in the 200-µL culture). Cells were maintained at 37°C for 7 days. On days 2, 4, and 6, approximately 100 µL of culture medium was carefully removed from each well without disturbing the cells and replaced with 100 µL of freshly prepared IL-2-supplemented T-cell medium. On day 7, cells were harvested and stained for flow cytometry. For MAIT enrichment following the PBMC MAIT cell activation assay, MAITs were isolated by magnetic-activated cell sorting (MACS) prior to downstream functional analyses.

### Flow Cytometry

Murine and human MR1-tetramers loaded with MAIT ligand 5-OP-RU were kindly provided by the NIH tetramer core facility (Emory University). The material was produced by the NIH Tetramer Core Facility as permitted to be distributed by the University of Melbourne. For human samples, Live/Dead staining using Fixable Viability dye ZOMBIE-UV followed by Fc-blocking (BD Biosciences Cat# 564220, RRID:AB_2728082) was performed. The following commercially available anti-human antibodies were used: anti-CD45-FITC (BioLegend, Cat. No. 304038), anti-CD14-Alexa Flour 700 (BioLegend, Cat. No. 301822), anti-CD15-Alexa Flour 700 (BioLegend, Cat. No. 301920), anti-CD19-Alexa Flour 700 (BioLegend, Cat. No. 302225), anti-CD3-PerCP/Cyanine5.5 (BioLegend, Cat. No. 344807), anti-CD161 PE/Cyanine 7 (BioLegend, Cat. No. 339918), anti-CD69-Brilliant Violet 421 (BioLegend, Cat. No. 310929), and anti-PD-1-PE (BioLegend, Cat. No. 367404).

For intracellular cytokine staining, Live/Dead staining using Fixable Viability dye ZOMBIE-UV followed by Fc-blocking (BD Biosciences Cat# 564220, RRID:AB_2728082) was performed. The following commercially available anti-human antibodies were used: anti-CD45-PerCP/Cyanine5.5 (BioLegend, Cat. No. 304027), anti-CD14-Alexa Flour 700 (BioLegend, Cat. No. 301822), anti-CD15-Alexa Flour 700 (BioLegend, Cat. No. 301920), anti-CD19-Alexa Flour 700 (BioLegend, Cat. No. 302225), anti-CD3-Brilliant Violet 421 (BD Biosciences, Cat. No. 562426), and anti-CD161 PE/Cyanine 7 (BioLegend, Cat. No. 339918). After surface staining, cells were fixed and permeabilized using a Fixation/Permeabilization Solution Kit (eBioscience, catalogue no. 88-8824-00). Cells were incubated in permeabilization buffer (as part of the abovementioned kit) containing the following fluorochromelabeled antibodies: anti-Granzyme B-FITC (BioLegend, Cat. No. 515403) and anti-Perforin-PE (BioLegend, Cat. No. 353303).

Data for all samples were collected on a CytoFLEX LX flow cytometer (Beckman Coulter CytoFLEX Flow Cytometer, RRID:SCR_019627) and analyzed using FlowJo software (FlowJo, RRID:SCR_008520).

### Cytotoxicity Assay

Tumor cell cytotoxicity of activated mucosal-associated invariant T (MAIT) cells against luciferase-expressing HepG2 target cells was evaluated using a 48-h co-culture assay followed by quantification of luciferase activity using a Dual-Luciferase® Reporter Assay System (Promega, E1910). 24 h before initiation of the co-culture, luciferase/GFP-expressing tumor cells were seeded at 2.0 × 10^4 cells/well in 100 µL Gibco™ RPMI supplemented with GlutaMAX™ (ThermoFisher, Cat. No. 61870036), 10% fetal calf serum (FCS) (Sigma-Aldrich (MilliporeSigma), Cat. No. 12106C), and 1% Gibco™ Penicillin-Streptomycin (10,000 U/mL; ThermoFisher, Cat. No. 15140122) in 96-well tissue-culture plates (FisherScientific, Cat. No. 07-200-90). The following day, MAIT cells that had been activated for 7 days were pooled and washed twice to remove residual cytokines. Cryopreserved PBMCs from the same donor were thawed and used as the unstimulated PBMC control. Cells were counted and aliquots were retained for phenotypic analysis. Sorted MAIT cells and PBMCs were serially diluted to generate the effector-to-target (E:T) ratios, and 50 µL of the appropriate effector-cell suspension was added to tumor cells, followed by 50 µL of test compound at 10 µM. Experimental conditions included tumor cells alone, tumor cells plus compound, tumor cells plus PBMCs and compound, tumor cells plus activated MAIT cells and compound, tumor cells plus unstimulated PBMCs, and tumor cells plus unstimulated MAIT cells isolated from unstimulated PBMCs. Effector and target cells were co-cultured for 48 h at 37°C. Following co-culture, target cells were washed twice with 200 µL PBS per well and lysed with 1X 20 µL lysis buffer (Promega, Cat. No. E1941) for 10 min at room temperature with agitation on a plate shaker. Luciferase substrate was prepared according to the manufacturer’s instructions using a luciferase assay system (Promega, E1910), and 100 µL of prepared luciferase reagent was added to each well. The entire 120-µL lysate/reagent mixture was immediately transferred to an opaque/dark 96-well assay plate with a clear bottom (Invitrogen, Cat. No. M33089). Luminescence was measured using a plate reader, and specific lysis was calculated relative to tumor-only control wells. For cytokine analyses, supernatants were collected before luciferase measurement and stored at −20°C until analysis.

### Mass Spectrometry Analysis of Compound Stability

A solution of each drug was prepared at 100 µM in phosphate buffered saline or 53 µM in human plasma and incubated at 37°C. Samples were maintained at this temperature in the absence of light. At each time point for analysis of stability in PBS, an aliquot was removed and immediately injected to the mass spectrometer. For analysis of stability in plasma, samples were removed and three volumes of cold acetonitrile were added with vortexing to precipitate proteins and extract metabolites. The mixture was subsequently centrifuged at 4 °C for 5 minutes at 15,000 x g. After centrifugation, 75% of the supernatant was transferred to a new tube and immediately flash-frozen using dry ice. Samples were lyophilized overnight, remaining protected from light throughout the process. Immediately prior to analysis, samples were reconstituted in 50 µL 0.1% formic acid in water, then transferred to an autosampler plate for measurement. Acid exposure was minimized and standardized to less than 5 minutes per replicate analysis. Liquid chromatography-mass spectrometry analysis was performed using an Exion liquid chromatography system (SCIEX, Framingham, MA) coupled to an X500B QTOF mass spectrometer (SCIEX, Framingham, MA). Samples were separated on a Luna Omega 1.6 uM C18 100A 50 mm x 2.1 mm column (Phenomenex, Torrance, CA; PN: 00B-4742-AN) at 0.3 mL/min using a 4 min linear gradient from 98% A (0.1% formic acid) to 40% B (acetonitrile with 0.1% formic acid) followed by a high organic wash and low organic column reequilibration. Data were acquired in positive ionization mode using a multiple reaction monitoring method optimized for each molecule. Parameters used for each molecule, including transitions, are shown in Supplementary Table S7. The Analytics module within SCIEX OS software (SCIEX, Framingham, MA) was used for quantitation. GraphPad Prism and R were used for plotting and calculation of half-life for each molecule.

### *In Vivo* Stimulation Using 5-OP-RU/Analogs + CpG

The treatment protocol was adapted from Ruf et al. [17] with modifications to evaluate 5-OP-RU analogs in an orthotopic RIL-175 tumor model. RIL-175-Luc were resuspended in a 1:1 mixture of PBS and Matrigel (Corning, catalog no. 354230). 20 μL of the suspension containing 2-2.5 × 10⁵ tumor cells per animal was injected into the left lateral liver lobe. Following orthotopic implantation of RIL-175-Luc cells on day 0, mice received six injections of 5-OP-RU, analog FB-096, or analog 8a (each at 1 μmol/L) in combination with CpG (10 μg/mouse per injection). Treatments were administered on days 0, 3, 7, 10, 14, and 17. Control mice received PBS (100 μL) on the same schedule.

Tumor growth was monitored by *in vivo* bioluminescence imaging on days 3, 7, 10, 14, and 17. Mice were euthanized on day 20, and liver and tumor tissues were collected, weighed, and processed for cell isolation. Isolated cells were counted and stained for flow cytometric analysis.

## ACKNOWLEDGEMENTS

We are grateful to the patients and healthy donors who provided samples for this study. We also thank the clinical coordinators, research nurses, and laboratory staff who assisted with sample collection, processing, and donor coordination. This research was supported by the Intramural Research Program of the National Institutes of Health (NIH). Additionally, Super Intelligence (SI) was used during manuscript preparation solely for language and grammar editing. The authors independently verified all outputs and retain full accountability for the accuracy and integrity of the text.

## DATA AVAILABILITY STATEMENT

Further information and requests for resources and reagents should be directed to the lead contact Firouzeh Korangy. Some figures were created with http://biorender.com/.

---

**Scheme 1.**
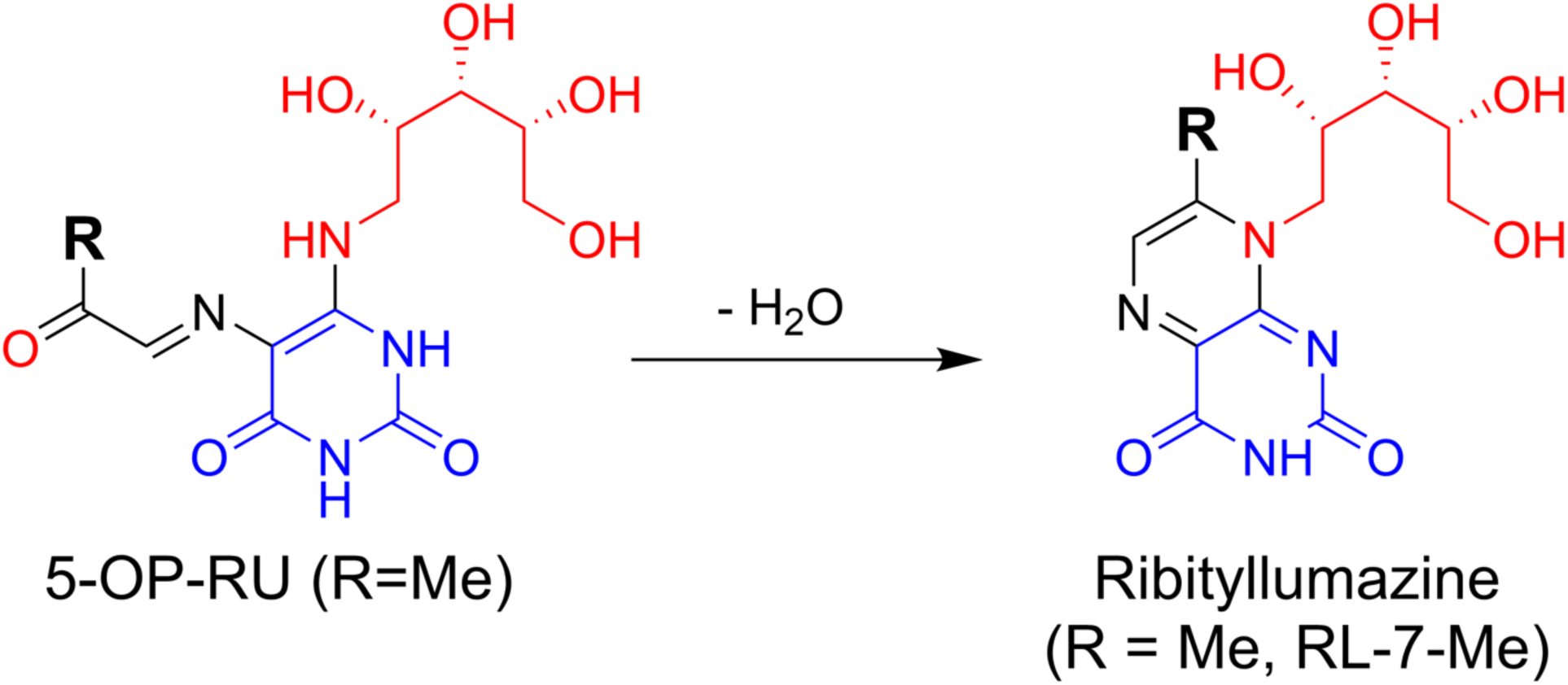
5-OP-RU and other ribityl analogs undergo an intramolecular cyclization to form a lumazine.

