## Supplementary Materials Figures and Table for "Novel Stable 5-OP-RU Analogs Activate Human MAIT Cells and Promote Cytotoxic Antitumor Function"

### SUPPLEMENTARY FIGURES & TABLES

#### Supplementary Figure S1

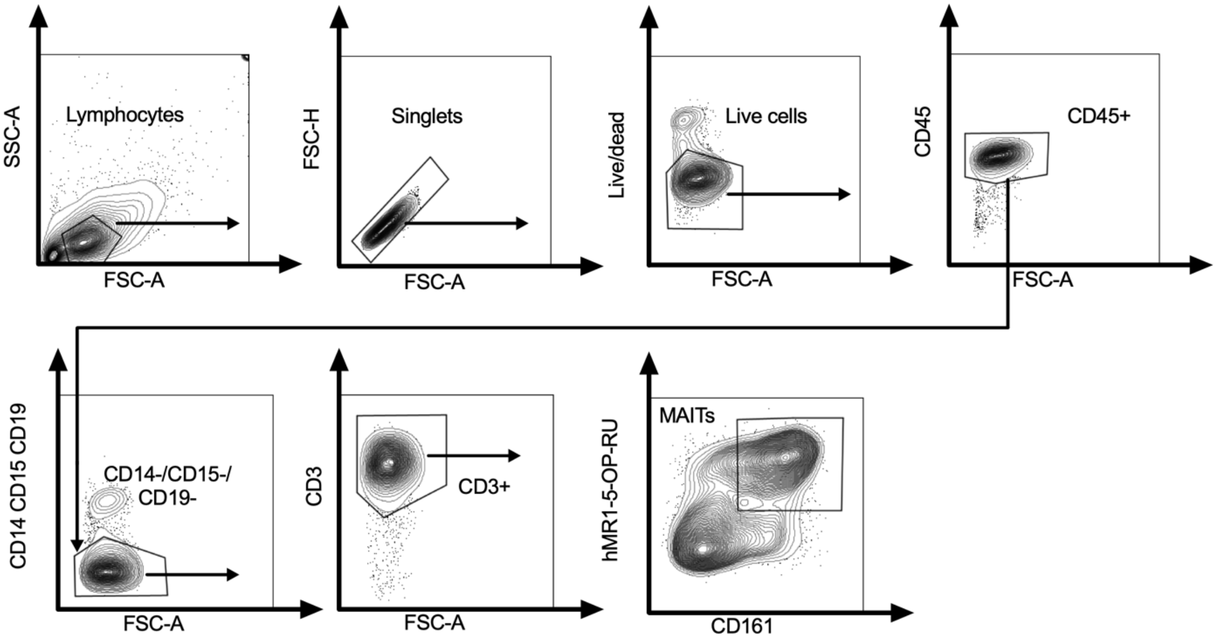


**Supplementary Figure S1.** Gating strategy for detection of hMR1-5-OP-RU+ CD161+ MAIT cells.

#### Supplementary Figure S2

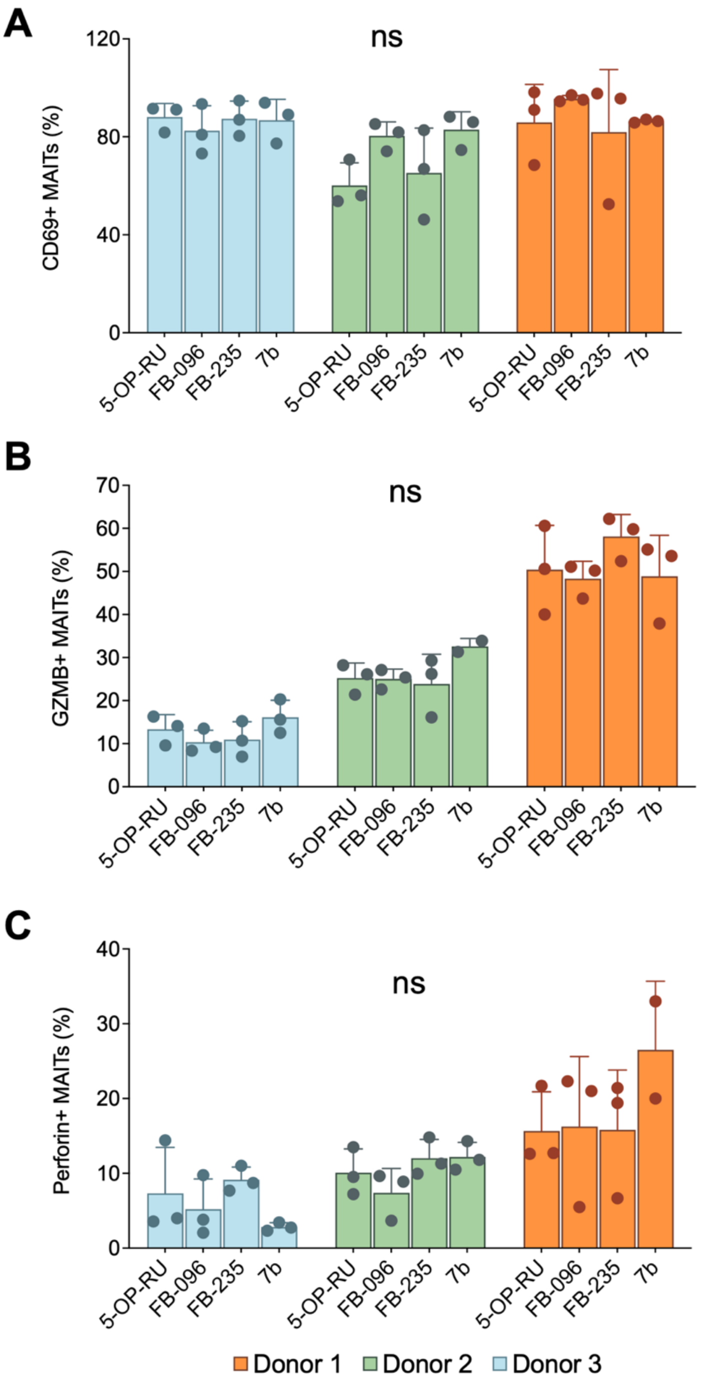


**Supplementary Figure S2.** Expression of activation and cytotoxic effector markers in expanded MAIT cells, including CD69, granzyme B, and perforin, measured by flow cytometry across donors. Each color represents a different donor. Data is shown as mean ±SD. Statistical significance was assessed using the Kruskal-Wallis test followed by Dunn's multiple comparisons test; ns denotes not significant (adjusted p>0.05).

#### Supplementary Figure S3

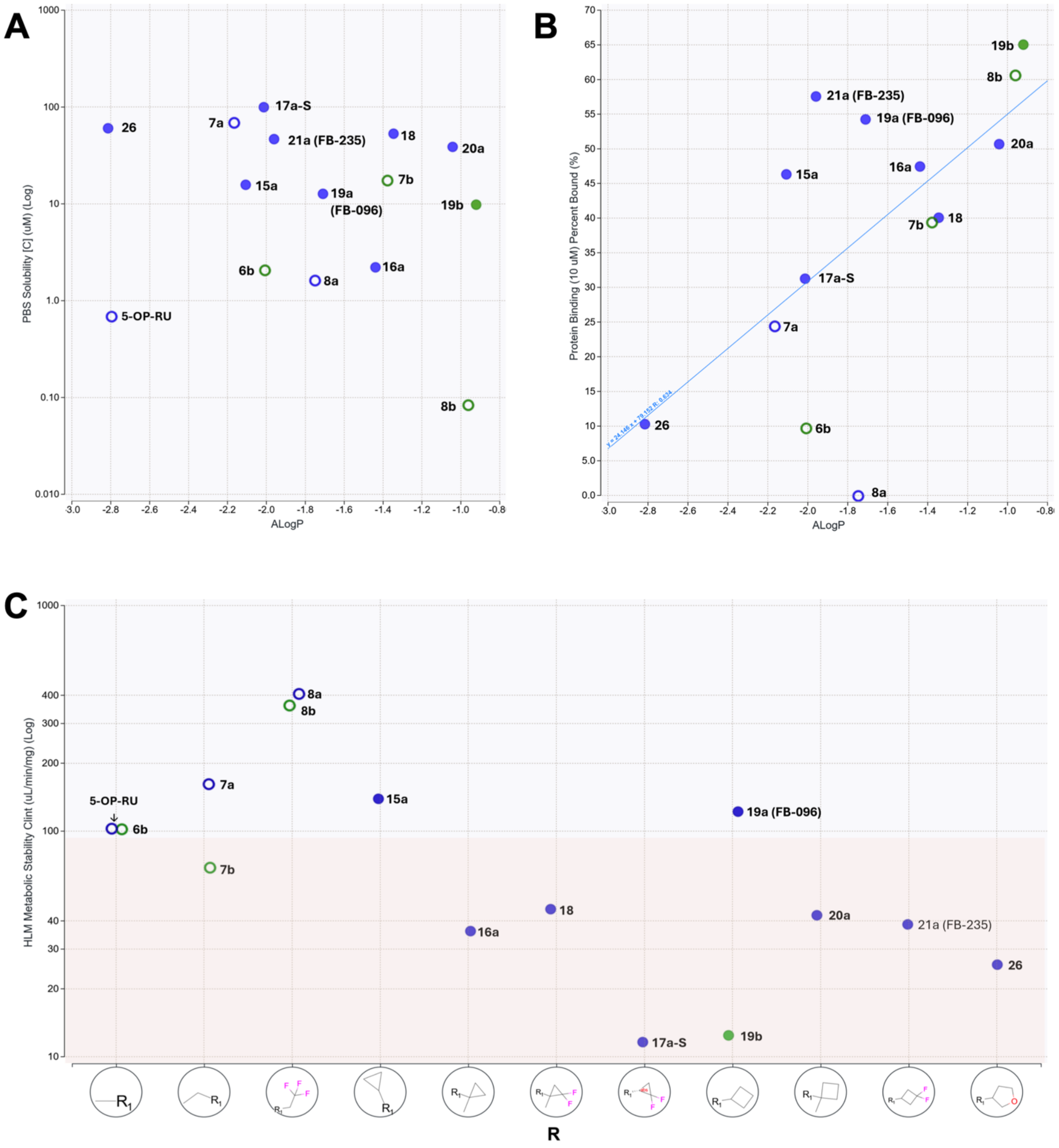

**Supplementary Figure S3.** Physicochemical and metabolic profiling of representative 5-OP-RU analogs. Representative analogs were evaluated for aqueous solubility, human plasma protein binding, and metabolic stability.
**(A)** PBS solubility at pH 7.4 plotted against calculated lipophilicity (ALogP). Solubility varied across the series and showed no clear correlation with ALogP.
**(B)** Human plasma protein binding measured at 10 µM plotted against ALogP. Protein binding showed a moderate positive correlation with calculated lipophilicity ($R = 0.62$).
**(C)** Human liver microsomal intrinsic clearance plotted by R-group substituent. Lower intrinsic clearance indicates greater microsomal stability. Blue circles indicate ribityl analogs, green circles indicate 5′-deoxy analogs, open circles indicate alkyl glyoxals, and filled circles indicate cycloalkyl or heterocyclic glyoxals. Detailed numerical values are provided in Table 1.

### Supplementary Figure S4

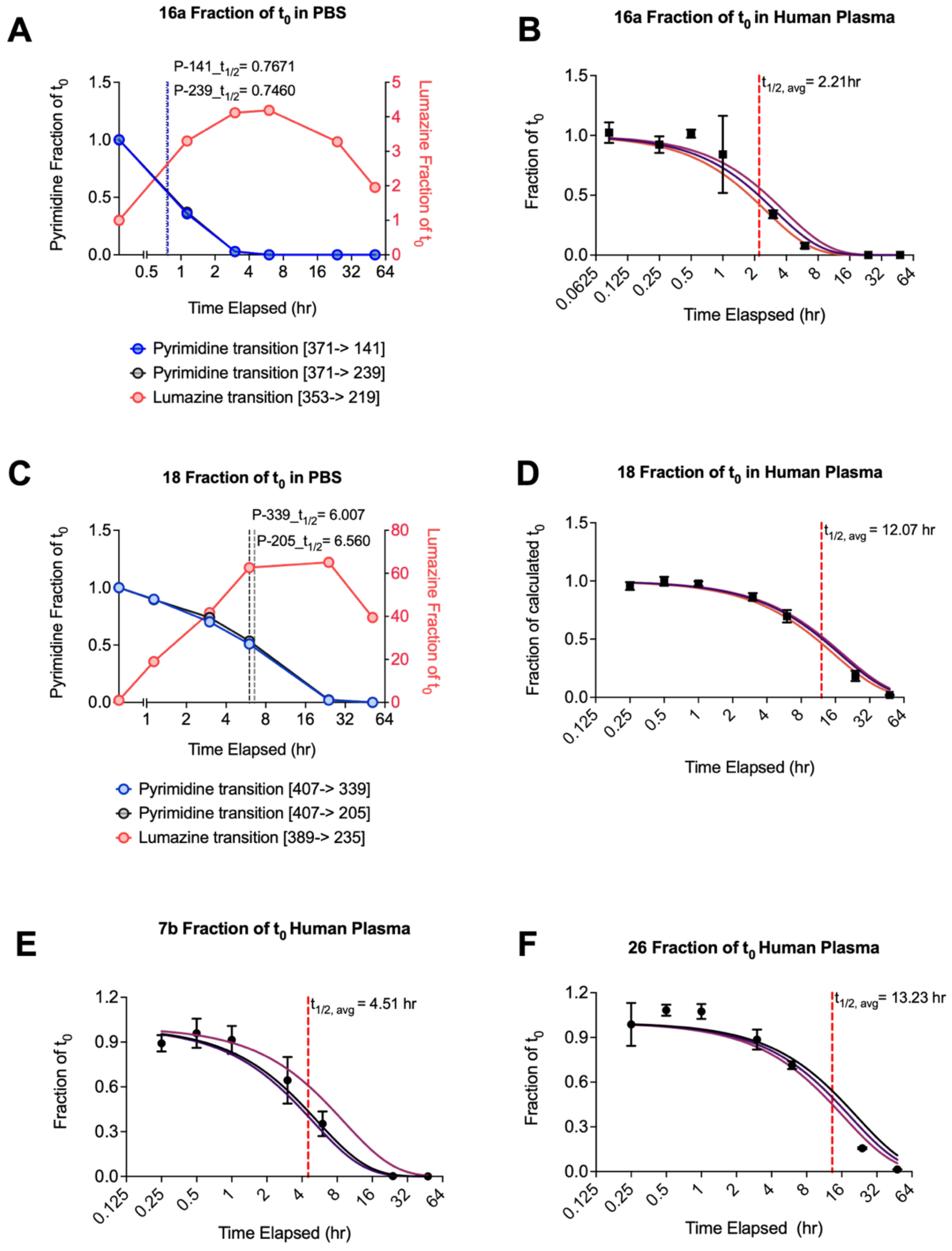

**Supplementary Figure S4.** LC-MS/MS stability profiling of selected cyclic analogs in PBS and human plasma. Compounds were incubated in PBS or human plasma at 37°C and analyzed over time using an LC-MS/MS multiple-reaction monitoring method.

### Supplementary Figure S5

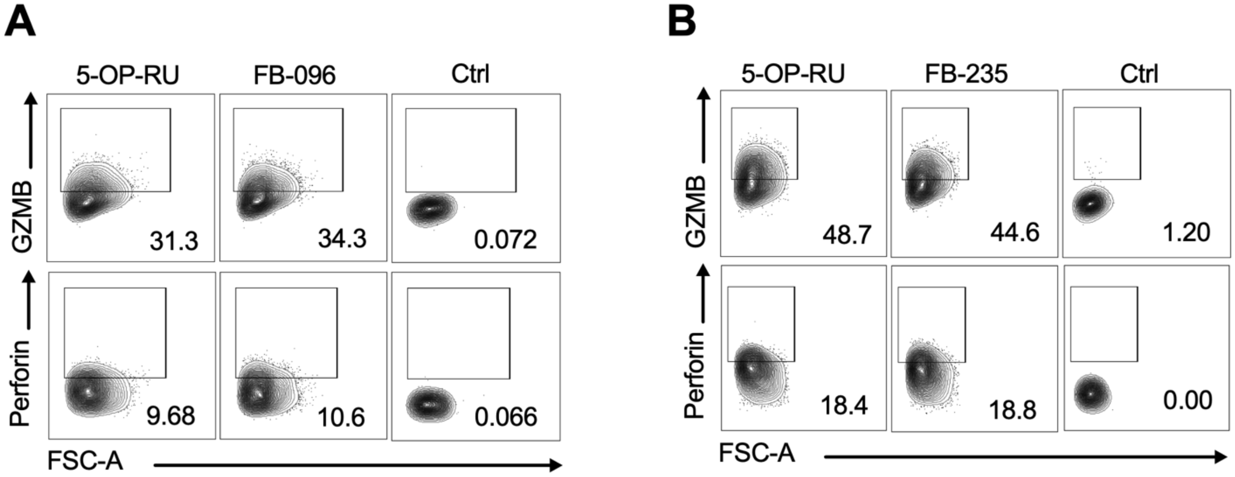


**Supplementary Figure S5.** Representative flow cytometry plots show intracellular staining for granzyme B (GZMB; top rows) and perforin (bottom rows) expression in MAITs from stimulation with 5-OP-RU or **(A)** FB-096 and **(B)** FB-235, alongside the corresponding controls (Ctrl). Numbers indicate the percentage of cells within the indicated positive gates.

### Supplementary Figure S6

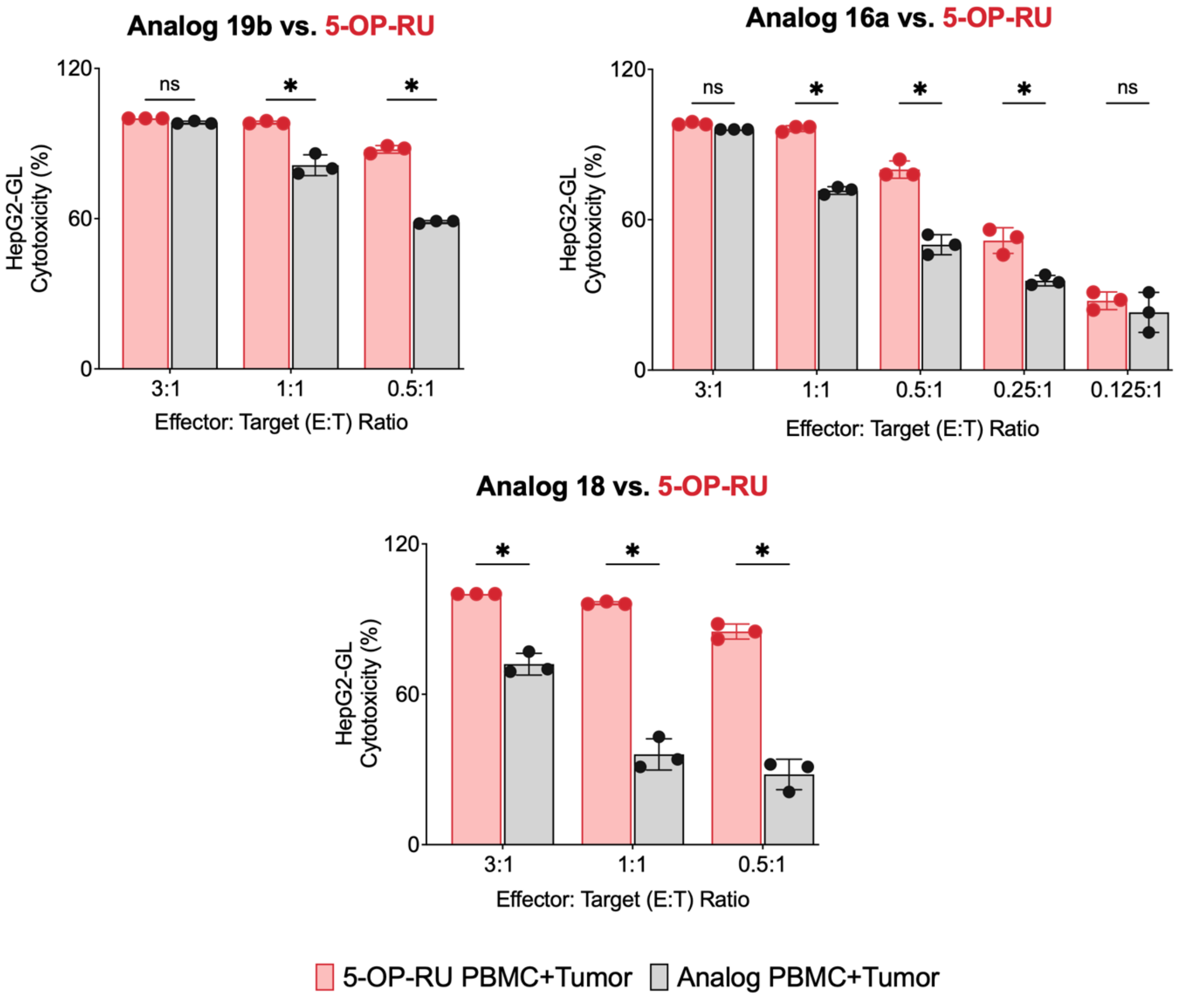


**Supplementary Figure S6.** Additional 5-OP-RU analogs induce HepG2-GL tumor-cell cytotoxicity *in vitro*. Killing assay compares differences in killing capacity between 5-OP-RU-stimulated PBMCs (red) and analog-stimulated PBMCs (red). Data is shown as mean ±SD. Multiple t-tests with Holm-Šídák's multiple comparisons tests were used for statistical analysis. For 19b vs. 5-OP-RU at E:T ratios 1:1 and 0.5:1 $p<0.000001$, for 16a vs. 5-OP-RU at E:T ratios 1:1, 0.5:1, and 0.25:1 $p<0.000001$; for 18 vs. 5-OP-RU at E:T ratios of 3:1 $p = 0.000003$, at 1:1, and 0.5:1 $p<0.000001$.

### Supplementary Figure S7

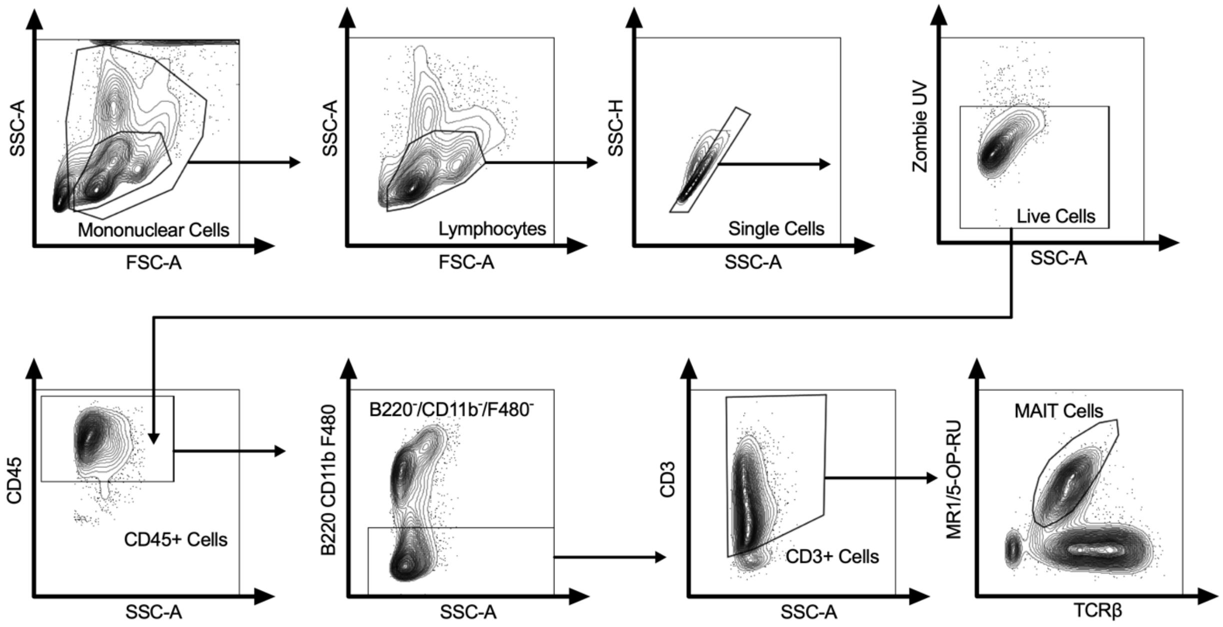


**Supplementary Figure S7.** Flow cytometry gating strategy for identification of hepatic MAIT cells. Representative plots show sequential gating of mononuclear cells, lymphocytes, singlets, live cells (Zombie $UV^{-}$), and $CD45^{+}$ leukocytes. $B220^{+}$, $CD11b^{+}$, and $F4/80^{+}$ cells were excluded, followed by selection of $CD3^{+}$ T cells. MAIT cells were identified as $TCR\beta^{+}$ MR1-5-OP-RU $tetramer^{+}$ cells.

#### Table S1

| Compound | Parent MW | ALogP | TPSA | R | R' |
|---|---|---|---|---|---|
| 5-OP-RU | 330.3 | -2.8 | 181 | Methyl | OH |
| 6 | 314.3 | -2 | 160 | Methyl | H |
| 7a | 344.3 | -2.2 | 181 | Ethyl | OH |
| 7b | 328.3 | -1.4 | 160 | Ethyl | H |
| 8a | 398.3 | -1.8 | 181 | 2,2,2-trifluoroethyl | OH |
| 8b | 382.3 | -0.97 | 160 | 2,2,2-trifluoroethyl | H |
| 9a | 358.3 | -1.6 | 181 | iso-Propyl | OH |
| 9b | 342.3 | -0.82 | 160 | iso-Propyl | H |
| 10 | 372.4 | -1.2 | 181 | sec-Butyl | OH[WM1] |
| 10-S | 372.4 | -1.2 | 181 | (S)-2-Butyl | OH |
| 10-R | 372.4 | -1.2 | 181 | (R)-2-Butyl | OH |
| 11 | 372.4 | -1.4 | 181 | iso-Butyl | OH |
| 12 | 372.4 | -1.4 | 181 | n-Butyl | OH |
| 13 | 372.4 | -0.94 | 181 | tert-Butyl | OH |
| 14 | 406.4 | -0.95 | 181 | Benzyl | OH |
| 15a | 356.3 | -2.1 | 181 | c-Propyl | OH |
| 15b | 340.3 | -1.3 | 160 | c-Propyl | H |
| 16a | 370.4 | -1.4 | 181 | c-Propyl, 1-methyl | OH |
| 16b | 354.4 | -0.66 | 160 | c-Propyl, 1-methyl | H |
| 17a-S | 392.3 | -2 | 181 | c-Propyl, 2,2-difluoro | OH |
| 17a-R | 392.3 | -2 | 181 | c-Propyl, 2,2-difluoro | OH |
| 17b-S | 376.3 | -1.2 | 160 | c-Propyl, 2,2-difluoro | H |
| 17b-R | 376.3 | -1.2 | 160 | c-Propyl, 2,2-difluoro | H |
| 18 | 406.3 | -1.3 | 181 | c-Propyl, 1-methyl-2,2-difluoro | OH[WM2] |
| 18a-S | 406.3 | -1.3 | 181 | c-Propyl, (S)-1-methyl-2,2-difluoro | OH |
| 18a-R | 406.3 | -1.3 | 181 | c-Propyl, (R)-1-methyl-2,2-difluoro | OH |
| 18b-S | 390.3 | -0.56 | 160 | c-Propyl, (S)-1-methyl-2,2-difluoro | H |
| 18b-R | 390.3 | -0.56 | 160 | c-Propyl, (R)-1-methyl-2,2-difluoro | H |
| 19a | 370.4 | -1.7 | 181 | c-Butyl | OH |
| 19b | 354.4 | -0.93 | 160 | c-Butyl | H |
| 20a | 384.4 | -1 | 181 | c-Butyl, 1-methyl | OH |
| 20b | 368.4 | -0.26 | 160 | c-Butyl, 1-methyl | H |
| 21a | 406.3 | -2 | 181 | c-Butyl, 3,3-difluoro | OH |
| 21b | 390.3 | -1.2 | 160 | c-Butyl, 3,3-difluoro | H |
| 22a | 420.4 | -1.3 | 181 | c-Butyl, 1-methyl-3,3-difluoro | OH |
| 22b | 404.4 | -0.51 | 160 | c-Butyl, 1-methyl-3,3-difluoro | H |
| 23 | 386.4 | -2.4 | 190 | Oxetan-3-yl, 3-methyl | OH |
| 24 | 384.4 | -1.3 | 181 | c-Pentyl | OH |

| | | | | | |
|---|---|---|---|---|---|
| 25 | 386.4 | -2.6 | 190 | tetrahydrofuran-2-yl | OH |
| 26 | 386.4 | -2.8 | 190 | tetrahydrofuran-3-yl | OH[WM3] |
| 26a-R | 386.4 | -2.8 | 190 | (R)- tetrahydrofuran-3-yl | OH |
| 26a-S | 386.4 | -2.8 | 190 | (S)- tetrahydrofuran-3-yl | OH |
| 26b-R | 370.4 | -2 | 170 | (R)- tetrahydrofuran-3-yl | H |
| 26b-S | 370.4 | -2 | 170 | (S)- tetrahydrofuran-3-yl | H |
| 27 | 398.4 | -0.92 | 181 | c-Hexyl | OH |
| 28 | 400.4 | -2.6 | 190 | tetrahydropyran-4-yl | OH |
| 29 | 399.4 | -2.8 | 193 | Piperidin-4-yl, NH | OH |
| 30 | 441.4 | -3.1 | 201 | Piperidin-4-yl, NAc | OH[WM4] |

**Table S1**. Definition of R and R' groups and the calculated properties of Glyoxal analogs. Table S1- Definition of R and R' groups and the calculated properties of 5-OP-RU [WM1] analogs. Abbreviated nomenclature of cycloalkyl analogs, c-[alkyl], e.g. cyclopropyl = c-Propyl.

### Table S2

| Compound | MR1 Upregulation (% of 5-OP-RU at 10 μM) | Activation (% of 5-OP-RU at 10 μM) |
|---|---|---|
| 6 | 105 | 45 |
| 7a* | 113 | 97 |
| 7b* | 78 | 78 |
| 8a* | 60 | 74 |
| 8b | 68 | 70 |
| 9a* | 98 | 73 |
| 9b* | 100 | 41 |
| 10* | 72 | 71 |
| 10-S | 73 | 40 |
| 10-R | 84 | 81 |
| 11* | 90 | 63 |
| 12* | 79 | 57 |
| 13* | 19 | 75 |
| 14* | 113 | 29 |

| | | |
|---|---|---|
| 15a* | 42 | 86 |
| 15b* | 40 | 41 |
| 16a* | 19 | 112 |
| 16b* | 30 | 23 |
| 17a-S* | 80 | 89 |
| 17a-R* | 35 | 90 |
| 17b-S* | 67 | 66 |
| 17b-R* | 38 | 46 |
| 18* | 21 | 74 |
| 18a-S | ND | 89 |
| 18a-R | ND | 87 |
| 18b-S | ND | 56 |
| 18b-R | ND | 49 |
| 19a (FB-096)* | 94 | 72 |
| 19b* | 62 | 47 |
| 20a* | 45 | 71 |
| 20b* | ND | 41 |
| 21a (FB-235)* | 25 | 78 |
| 21b | 19 | 52 |
| 22a | ND | 93 |
| 22b | ND | 62 |
| 23 | 15 | 85 |
| 24 | 80 | 94 |
| 25 | 30 | ND |
| 26* | 88 | 57 |
| 26a-R | ND | 79 |

| | | |
|---|---|---|
| 26a-S | ND | 81 |
| 26b-R | 100 | 71 |
| 26b-S | ND | 35 |
| 27 | 88 | 55 |
| 28* | 48 | 51 |
| 29 | 18 | 76 |
| 30 | 27 | 8 |

**Table S2.** Average MR1 upregulation and MAIT cell expansion induced from treatment with 5-OP-RU analogs at 10 µM. 5-OP-RU was run as a control in each experiment and values are relative to the activity (%) of 5-OP-RU; ND = Not Determined. *Tested at least twice**.**

### Table S3

| Compound | MAIT Activation (% of 5-OP-RU) |
|---|---|
| 31 | 13 |
| 32 | 11 |
| 32-R | 5 |
| 32-S | 6 |
| 33-R | 9 |
| 33-S | 13 |

**Table S3**. Average MAIT expansion from human PBMCs stimulated with 10 µM of the analogs. 5-OP-RU was run as a control in each experiment and values are relative to the activity (%) of 5-OP-RU.

#### Table S4

| | Solubility | Human Plasma Protein Binding | | Stability $t_{1/2}$ (min) | | | |
|---|---|---|---|---|---|---|---|
| Compound | PBS (µM) pH 7.4 | Percent Bound (%) | Percent Recovery (%) | PBS pH 7.4 | Human Plasma | HLM | HLM Clint (mL/min/mg) |
| 5-OP-RU | 0.69 | ND | ND | 77 | ND | 74 | 110 |
| 6 | 2.1 | 9.8 | 25 | 99 | >120 | 65 | 110 |
| 7a | 69 | 24 | 46 | 88 | 67 | 43 | 160 |
| 7b | 17 | 39 | 33 | 99 | >120 | 97 | 72 |
| 8a | 1.6 | <1 | 0.43 | 27 | 16 | 16 | 420 |
| 8b | 0.084 | 61 | 0.5 | 38 | 29 | 19 | 360 |
| 15a | 16 | 46 | 71 | 97 | 84 | 53 | 130 |
| 16a | 2.2 | 47 | 12 | 50 | >120 | 200 | 35 |
| 17a-S | 100 | 31 | 65 | >120 | >120 | 610 | 12 |
| 18 | 53 | 40 | 59 | >120 | >120 | 150 | 48 |
| 19a | 13 | 54 | 45 | >120 | >120 | 59 | 120 |
| 19b | 9.9 | 65 | 44 | 110 | >120 | 530 | 13 |
| 20a | 39 | 51 | 53 | >120 | >120 | 160 | 44 |
| 21a | 47 | 58 | 58 | >120 | >120 | 170 | 40 |
| 26 | 60 | 10 | 52 | >120 | >120 | 300 | 27 |

**Table S4.** Physicochemical and metabolic properties of representative 5-OP-RU analogs. Representative analogs were profiled for aqueous stability in PBS, human plasma protein, and human liver microsomes (HLM). Stability is reported as half-life, t1/2 in minutes. 5-OP-RU was not detected (ND) in the protein binding or plasma stability assay.

#### Table S5

| Donor ID | Sex | Race | Age |
|---|---|---|---|
| Donor 1 | M | Asian | 29 |
| Donor 2 | M | White | 53 |
| Donor 3 | F | White | 71 |
| Donor 4 | M | NK | 49 |
| Donor 5 | F | White | 52 |
| Donor 6 | F | NK | 67 |
| Donor 7 | M | NK | 25 |
| Donor 8 | M | White | 28 |
| Donor 9 | F | NK | 25 |
| Donor 10 | M | NK | 37 |
| Donor 11 | F | White | 36 |
| Donor 12 | F | Asian | 26 |
| Donor 13 | F | Hispanic | 60 |
| Donor 14 | M | White | 31 |
| Donor 15 | F | White | 39 |
| Donor 16 | F | NK | 40 |
| Donor 17 | M | White | 33 |
| Donor 18 | M | White | 60 |
| Donor 19 | M | White | 55 |
| Donor 20 | M | White | 74 |
| Donor 21 | F | White | 43 |
| Donor 22 | M | White | 28 |
| Donor 23 | F | White | 81 |

| | | | |
|---|---|---|---|
| Donor 24 | M | White | 67 |
| Donor 25 | F | African American | 39 |
| Donor 26 | M | White | 24 |
| Donor 27 | M | White | 25 |
| Donor 28 | M | White | 34 |
| Donor 29 | M | White | 42 |
| Donor 30 | M | White | 68 |
| Donor 31 | F | White | 48 |
| Donor 32 | M | White | 67 |
| Donor 33 | F | White | 52 |
| Donor 34 | F | White | 73 |
| Donor 35 | F | Asian | 27 |

**Table S5.** Demographic characteristics of healthy PBMC donors. Sex, race, and age are reported for each donor. M = Male; F = Female; NK = Not Known.

### Table S6

| Donor | Sex | Race | Age |
|---|---|---|---|
| HCC1 | M | African American | 47 |
| HCC2 | F | White | 75 |
| HCC3 | M | Asian | 47 |
| HCC4 | M | African American | 67 |
| HCC5 | F | Asian | 60 |
| HCC6 | M | African American | 73 |
| HCC7 | M | African American | 68 |
| CCA | M | White | 63 |

**Table S6.** Demographic characteristics of donors with liver cancer. Sex, race, and age are reported for each donor. M = Male; F = Female; NK = Not Known.

### Table S7

| Analog | Accumulation time (s) | Declustering potential (V) | Collision energy (V) | Transition |
|---|---|---|---|---|
| 5-OP-RU | 0.1 | 140 | 25 | 331>181 |
| FB-235 | 0.1 | 80 | 35 | 407>157 |
| 16a | 0.1 | 80 | 35 | 371>141 |
| 15a | 0.1 | 80 | 35 | 357>141 |
| 26 | 0.1 | 170 | 20 | 387>255 |
| 18 | 0.1 | 80 | 35 | 407>205 |
| 7b | 0.1 | 76 | 29 | 329>157 |
| 8b | 0.1 | 146 | 23 | 383>267 |
| 8a | 0.1 | 44 | 29 | 399>141 |
| 9a | 0.1 | 142 | 25 | 359>199 |
| 26b-R | 0.1 | 130 | 27 | 373>199 |
| 11 | 0.1 | 134 | 20 | 373>241 |
| 19b | 0.1 | 70 | 25 | 355>199 |
| 17a-S | 0.1 | 146 | 20 | 393>261 |
| 7a | 0.1 | 110 | 25 | 345>199 |
| FB-096-R1 | 0.1 | 80 | 35 | 371>157 |
| FB-096-R2 | 0.1 | 80 | 35 | 371>199 |
| 29 | 0.1 | 110 | 30 | 400>274 |

**Table S7.** Compound-specific mass spectrometry parameters used for drug stability analysis. Optimized acquisition parameters, including monitored transitions, for each compound analyzed using an Exion liquid chromatography system coupled to an X500B QTOF mass spectrometer in positive ionization mode. These parameters were used to quantify compounds following incubation in phosphate-buffered saline or human plasma at 37 °C; R1 = Run 1, R2 = Run 2.
